# Assessing intra-lab precision and inter-lab repeatability of outgrowth assays of HIV-1 latent reservoir size

**DOI:** 10.1101/329672

**Authors:** Daniel I. S. Rosenbloom, Peter Bacchetti, Mars Stone, Xutao Deng, Ronald J. Bosch, Douglas D. Richman, Janet D. Siliciano, John W. Mellors, Steven G. Deeks, Roger G. Ptak, Rebecca Hoh, Sheila M. Keating, Melanie Dimapasoc, Marta Massanella, Jun Lai, Michele D. Sobolewski, Deanna A. Kulpa, Michael P. Busch, for the Reservoir Assay Validation and Evaluation Network (RAVEN) Study Group

**Affiliations:** Department of Systems Biology, Columbia University College of Physicians and Surgeons, New York, New York, United States of America; Department of Biomedical Informatics, Columbia University College of Physicians and Surgeons, New York, New York, United States of America; Department of Epidemiology and Biostatistics, University of California San Francisco, San Francisco, California, United States of America; Blood Systems Research Institute, San Francisco, California, United States of America; Center for Biostatistics in AIDS Research, Harvard T. H. Chan School of Public Health, Boston, Massachusetts, United States of America; University of California San Diego, La Jolla, California, United States of America; Veterans Affairs San Diego Healthcare System, San Diego, California, United States of America; Department of Medicine, Johns Hopkins University School of Medicine, Baltimore, Maryland, United States of America; Division of Infectious Diseases, Department of Medicine, University of Pittsburgh School of Medicine, Pittsburgh, Pennsylvania, United States of America; Division of HIV, Infectious Diseases and Global Medicine, University of California San Francisco, San Francisco, California, United States of America; Southern Research, Frederick, Maryland, United States of America; Department of Pediatrics, Emory University, Atlanta, Georgia, United States of America; Department of Laboratory Medicine, University of California, San Francisco, San Francisco, California, United States of America

## Abstract

Quantitative viral outgrowth assays (QVOA) use limiting dilutions of CD4+ T cells to measure the size of the latent HIV-1 reservoir, a major obstacle to curing HIV-1. Efforts to reduce the reservoir require assays that can reliably quantify its size in blood and tissues. Although QVOA is regarded as a “gold standard” for reservoir measurement, little is known about its accuracy and precision or about how cell storage conditions or laboratory-specific practices affect results. Owing to this lack of knowledge, confidence intervals around reservoir size estimates – as well as judgments of the ability of therapeutic interventions to alter the size of the replication-competent but transcriptionally inactive latent reservoir – rely on theoretical statistical assumptions about dilution assays. To address this gap, we have carried out a Bayesian statistical analysis of QVOA reliability on 75 split samples of peripheral blood mononuclear cells (PBMC) from 5 antiretroviral therapy (ART)-suppressed participants, measured using four different QVOAs at separate labs, estimating assay precision and the effect of frozen cell storage on estimated reservoir size. We found that typical assay results are expected to differ from the true value by a factor of 1.6 to 1.9 up or down. Systematic assay differences comprised a 24-fold range between the assays with highest and lowest scales, likely reflecting differences in viral outgrowth readout and input cell stimulation protocols. We also found that controlled-rate freezing and storage of samples did not cause substantial differences in QVOA compared to use of fresh cells (95% probability of < 2-fold change), supporting continued use of frozen storage to allow transport and batched analysis of samples. Finally, we simulated an early-phase clinical trial to demonstrate that batched analysis of pre- and post-therapy samples may increase power to detect a three-fold reservoir reduction by 15 to 24 percentage points.

**Author summary:** The latent reservoir of resting CD4^+^ T cells is a major, if not the primary, obstacle to curing HIV. Quantitative viral outgrowth assays (QVOAs) are used to measure the latent reservoir in ART-suppressed HIV-infected people. Using QVOA is difficult, however, as the fraction of cells constituting the latent reservoir is typically about one in one million, far lower than other infectious disease biomarkers. To study reliability of these assays, we distributed 75 PBMC samples from five ART-suppressed HIV-infected participants among four labs, each conducting QVOA and following prespecified sample batching procedures. Using a Bayesian statistical method, we analyzed detailed assay output to understand how results varied within batches, between batches, and between labs. We found that, if batch variation can be controlled (i.e., a lab assays all samples in one batch), typical assay results are expected to differ from the true value by a factor of 1.6 to 1.9 up or down. We also found that freezing, storing, and thawing samples for later analysis caused no more than a 2-fold change in results. These outcomes, and the statistical methods developed to obtain them, should lead towards more precise and powerful assessments of HIV cure strategies.

## Introduction

The latent HIV-1 reservoir that persists following treatment with suppressive ART exists primarily in resting CD4^+^ T cells and is an obstacle to eradicating HIV-1 [1–4]. There are substantial ongoing efforts to eliminate or reduce the size of this reservoir [5–7]. Evaluating such efforts requires assays that can reliably quantify its size in blood and tissues in order to monitor its changes during curative intervention strategies. Replication-competent HIV-1 can be measured by QVOA. These terminal dilution assays place known numbers of resting CD4^+^ T cells in culture wells, usually in serial dilutions of cells that cover several orders of magnitude, with replicate wells at each dilution. The CD4^+^ T cells are activated before co-culture with cells that are highly permissive for primary strains of HIV. The propagation of HIV replication is detected by an assay for either p24 antigen or HIV RNA in the supernatant of these co-cultures over a two-to three-week period [8]. Each well is read as negative or positive, which means that replication-competent virus was present in at least one of the cells in the well. The number of infectious units per million cells (IUPM) is then estimated by maximum likelihood assuming single-hit Poisson dynamics [9]. This approach has represented the “gold standard,” because it measures replication-competent virus in latently infected cells, which is crucial because the majority of integrated HIV-1 DNA is replication-defective [10–12].

Use of QVOA presents both practical and statistical challenges, many of which are attributable to the rarity of the target entities: often on the order of only one CD4^+^ T cell in a million is positive for replication-competent virus by a single-round QVOA. These challenges include: 1) QVOA requires that large volumes of blood be collected to generate large numbers of Ficoll-purified peripheral blood mononuclear cells (PBMC), which are generally further processed without freezing/thawing, into input resting CD4^+^ T cells, 2) each assay takes weeks to complete and is expensive (~$3,000), 3) substantial personnel time is required for cell purification, culture and monitoring supernatants for HIV replication, limiting test throughput to two to four QVOAs per lab per week, 4) not all replication-competent virus is detected by a single QVOA [11], and 5) different laboratories employ varying methods [8].

In addition, the performance characteristics of QVOAs performed within and between labs have not been carefully evaluated, a gap that this study was designed to address. The rarity of infectious units complicates the analysis of performance characteristics, because even split samples from the same collection can have large relative differences in the true numbers of infectious units that they contain. In addition, the number of infectious units in positive wells is not known, which also adds to the assays’ variability. We describe in the next section Markov-chain Monte Carlo (MCMC) methods that we developed to account for these inevitable background sources of variation while estimating additional variation, including batch effects and inter-lab variation, as well as assessing the impact of freezing PBMC samples on assay performance. We also describe simulations to validate these methods and to assess the implications of the parameter estimates. We then present the results of our method-validation simulations, estimated model parameters based on the results of four QVOAs applied to 75 split samples, and simulation results evaluating some implications of the models, before concluding with some additional discussion.

## Materials and methods

### Ethics Statement

Participants in the RAVEN project are enrolled and followed as part of the UCSF OPTIONS and SCOPE programs, with specific consent for apheresis collections and testing for this study as approved by the UCSF Committee on Human Research (IRB) # 10-03244.

### Experimental design

The five ART-suppressed HIV-1 infected participants for the current study were selected to have diverse replication-competent reservoirs based on QVOA results from a previously published study [10]. Leukapheresis collections from five HIV+ participants and one uninfected control participant were divided into 12 aliquots (control) or 15 aliquots (each HIV+ participant). Each aliquot contained roughly 300 – 750 million PBMC as requested by the testing labs. One aliquot from each HIV+ participant comprised the fresh panel; all other aliquots were stored at −180° C and comprised the frozen panel.

Four labs participated in the study: University of Pittsburgh (U. Pitt.), University of California San Diego (UCSD), Johns Hopkins University (JHU), and Southern Research (SR). One aliquot per participant from the fresh panel was distributed for immediate testing to three of the labs (all except SR). The frozen panel included 18 uniquely coded liquid nitrogen-cryopreserved PBMC aliquots that were distributed to all four labs for testing 4 – 12 months after freezing, depending on lab testing capacity. Aliquots were blinded as to participant and aliquot identity, except that all labs had knowledge that the negative control was not included in the fresh panel. Within each lab, the frozen panel was analyzed in balanced batches of two aliquots each, designed to enable measurement of both within-batch and between-batch variation (Figure 1).

**Fig 1.**
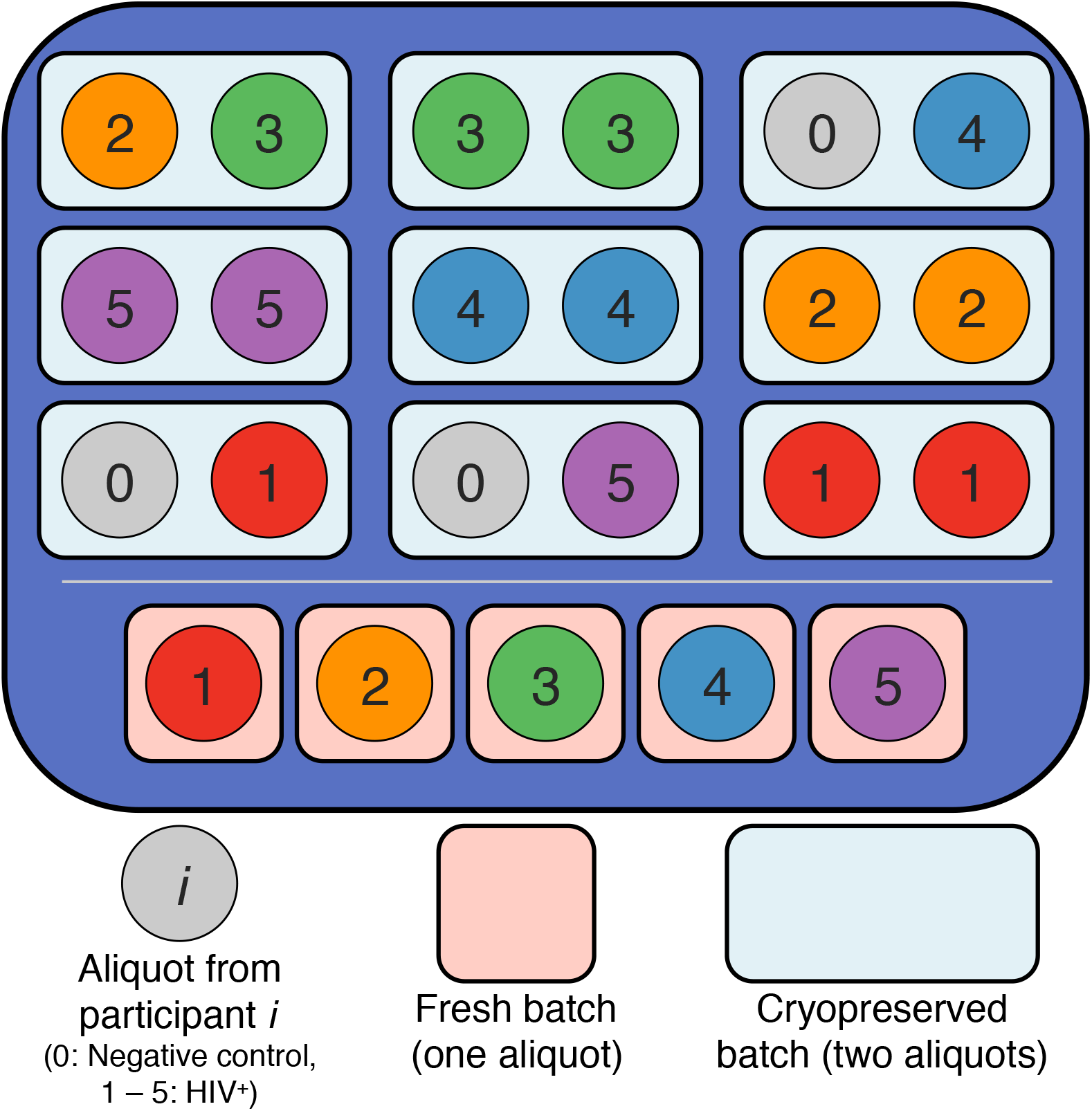
Experimental design at one of the four labs (U. Pitt.). *Fresh panel* (five batches): One aliquot from each HIV^+^ participant was studied fresh and was not batched with any other aliquots. *Frozen panel* (nine batches): Three aliquots from each participant were cryopreserved and batched together with one other aliquot. Five batches contain two aliquots from the same HIV^+^ participant. Two HIV^+^ participants are chosen to supply one aliquot to the same batch (here, participants 2 and 3). The remaining three batches contain an aliquot from one of the remaining HIV^+^ participants and an aliquot from the negative control. In each lab, different HIV^+^ participants are chosen for the mixed batch (see S1 Table for complete experimental design).

Labs thawed aliquots (if frozen), isolated CD4^+^ T cells (CD4s), and performed QVOA per lab protocol (S2 Table). S3 Table provides the well configurations (cell input counts and number of replicate wells) used for each aliquot. While QVOA output is typically reported as an estimated infection frequency and confidence interval (expressed as infectious units per million, or IUPM), all labs reported individual well outcomes (positive or negative for viral outgrowth). Reporting at this higher level of granularity allowed for more accurate statistical modeling. Individual laboratory reports were unblinded, checked for transcription errors if there were discrepancies in IUPMs calculated by the RAVEN statistical team and IUPMs reported by the laboratories, and then compiled for statistical analysis. S4 Table reports the resulting dataset.

The JHU lab used two different protocols for the fresh and frozen panels: For the fresh panel, viral outgrowth was measured at day 7 and 14 of coculture for all aliquots, but continuation to day 21 was contingent on the results at day 14. For the frozen panel, viral outgrowth was measured at days 7, 14 and 21 for all aliquots. Owing to this variance in methods, we present analysis of the data in two ways, differing in treatment of the JHU lab: Our primary analysis used cumulative QVOA results through day 21 measurements from the JHU frozen panel and no measurements from the JHU fresh panel, while our secondary analysis used day 14 measurements from both panels. As some wells require the full 21 days for outgrowth to be evident [13], the day 21 measurement is more sensitive, and so the primary analysis may yield more relevant characterization of QVOA precision. The secondary analysis, however, draws upon three labs instead of two for the fresh/frozen comparison and may yield more relevant characterization of the effect of cryopreservation. Unless otherwise stated, all experimental results reported draw upon the primary analysis.

### Analytical methods

As noted in the Introduction, some variability in measured IUPM is unavoidable even for a perfect assay, due to Poisson sampling variation and uncertainty about the number of infectious units that were present in positive wells. We therefore developed a statistical model that estimates additional sources of variation beyond this unavoidable background. The design permitted identification of extra variation at the aliquot level, batch-to-batch variation within each lab, and lab-to-lab variation. Below are details of the statistical model that accomplishes this task, along with our methods for fitting the model. We did not include the control participant in this modeling.

#### Statistical model

Wells in the experiment are indexed by study participant from which the sample was obtained (*i* = 1… 5), aliquot into which the sample was divided (*j* = 1… 15 for each participant, with 3 – 4 aliquots sent to each lab), assay batch (*k* = 1… 14 for each lab), and the lab where the assay was performed (*l* = 1 … 4, referring to U. Pitt., UCSD, JHU, and SR, respectively). Viral outgrowth was assumed to follow a Poisson model, in which a well containing *n* input cells has independent probability of being positive equal to

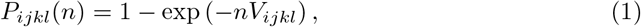

where *V_ijkl_* is the probability that a single cell in the well is capable of causing outgrowth,

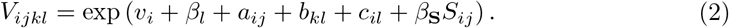

In this expression, *S_ij_* is set to one if aliquot *j* from participant *i* was cryopreserved before the assay, and otherwise is set to zero. Each other term represents a fixed or random effect (Table 1).

**Table 1.**
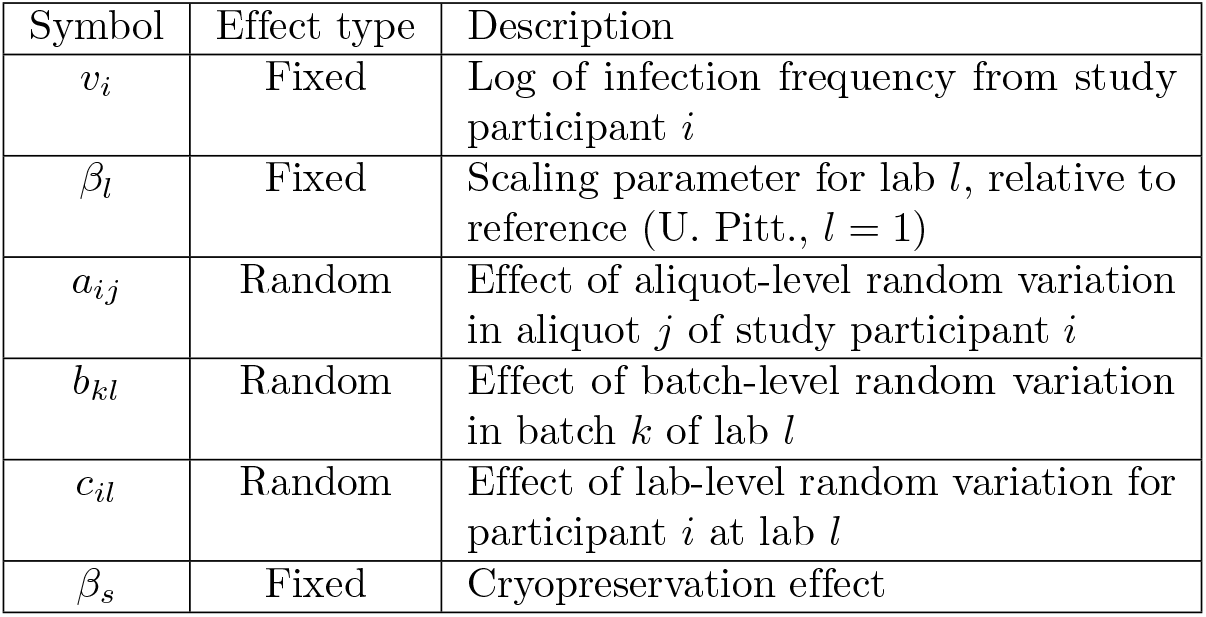
Fixed and random effects in the model of outgrowth, eq. (2)

The random effects in the model (2) are normal i.i.d. variates:

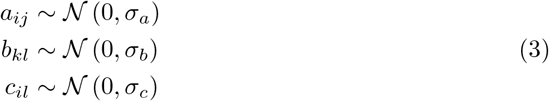

In this framework, *σ_a_* quantifies extra variation that is distinct for each single aliquot, beyond what would be expected from Poisson sampling variability and binary readout of wells. It may be thought of as an overdispersion parameter, similar to the extra-Poisson dispersion that is present in the negative binomial distribution. The parameter *σ_b_* reflects additional variation that equally influences both aliquots in the same batch, reflecting variation in laboratory personnel, condition of reagants used, or other aspects of laboratory environment. The random perturbations *c_il_* apply to all split samples from the same participant at a particular lab, but differ for the same participant at different labs. The parameter *σ_c_* can therefore be thought of as reflecting participant-specific differences in exactly what each lab measures, beyond the systematic scale differences *β_l_* that apply equally to all participants. A non-zero *σ_c_* indicates that each lab is measuring a quantity that correlates imperfectly with the true latent reservoir size, with smaller *σ_c_* corresponding to better correlation and therefore less variation between labs in what they are actually measuring.

#### Maximum likelihood estimation

We encountered difficulties fitting the above model by maximum likelihood, and therefore focused primarily on Bayesian estimation as described in the next section. We did not identify any existing maximum likelihood procedures in the software packages SAS, Stata, or R that would apply directly to the above model. We therefore considered a variation in which exp(*a_ij_*) follow a Gamma distribution, 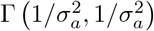. This results in the number of infectious units in an aliquot having a gamma mixture of Poisson distributions, which follows a negative binomial distribution with mean dispersion parameter 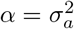 [14]. To fit the resulting model using mixed effects negative binomial models, we calculated for each aliquot stand-in data *r_ij_* and *N_ij_* such that a Poisson model applied to *r_ij_* with offset log(*N_ij_*) (or equivalently, including *N_ij_* as the “exposure” variable) results in the same estimated IUPM and confidence interval as the standard maximum likelihood calculations for the well-by-well data [9,15]. We then attempted to use software for mixed effects negative binomial regression (Stata v13 `menbreg` command, R x64 3.4.0 `glmer.nb` command in the `lme4` package v1.1-13, or SAS v9.4 `proc Glimmix`) to fit the needed model to the *r_ij_* and *N_ij_*, with crossed random effects *b_kl_* and *c_il_*.

#### Markov-chain Monte Carlo estimation

Given the difficulties with maximum likelihood estimation and the versatility of Markov-chain Monte Carlo (MCMC) approaches, we developed a Bayesian estimation framework. In order to obtain posterior distributions that mainly reflect the evidence provided by our data, we aimed to use prior distribution assumptions for the parameters that were “weak.” To make our assumptions neutral about the existence of nonzero random effects, we fit model versions with and without each of *σ_a_, σ_b_*, and *σ_c_*, with equal prior probability for each.

A uniform or normal distribution was used as prior for each fixed-effect parameter, and a half-Cauchy prior was used for each random effect parameter:

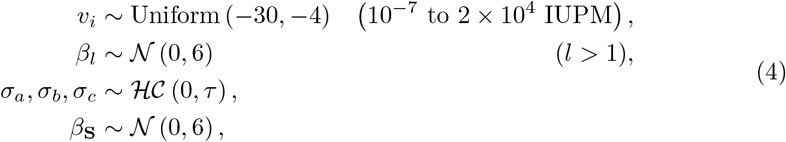

where Cauchy scale parameters *τ* = 1, 2, 3,4 were tried.

Eight versions of the model were fitted, one for each combination of the three random effects listed in Table 1. These were all given equal prior probability, making the effective prior for each random effect an equal mixture of the half-Cauchy noted above and a point mass at zero. Posterior distributions for parameters were estimated using PyStan 2.12 [16], with four parallel chains of 2500 iterations each. The first half of each chain was discarded as warmup. A posterior probability for each model was calculated using the Watanabe-Akaike Information Criterion (WAIC) [17]. Complete model specifications in Stan are provided (S1 Appendix).

A set of 1000 independent samples from each of the eight models’ posterior parameter distribution was obtained. Using model posterior probabilities as weights, weighted quantiles of these 8000 were computed. Posterior medians and 95% credible intervals (spanning 2.5 to 97.5 percentiles) were reported. Unless otherwise noted, all reported results use these ensemble estimates. Where a sample from the joint posterior was required, 1000 of the 8000 samples were selected, roughly proportional to the model weights (“complete ensemble”). An alternate ensemble of 1000 was also constructed setting batch effect to zero; in this case only four of the eight models were included (“batch variation-free ensemble”).

Most estimates are reported as fold-changes, that is, the exponential of a model parameter. Random variation at the aliquot and batch levels combined is reported as exp 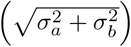; variation at all three levels combined is reported as 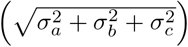.

#### Validation by simulation

For single-lab simulations ignoring cryopreservation, the five participants’ IUPMs were based on preliminary estimates from fresh samples: 0.104, 0.591, 0.654, 0.835, 1.454. Three levels of extra-Poisson variation were considered: *Large variation* (*σ_a_* = *σ_b_* = 0.7), *moderate variation* (*σ_a_* = *σ_b_* = 0.2), and no *variation* (*σ_a_* = *σ_b_* = 0). The simulated assay used six replicate wells of cell inputs 1,000,000, 300,000, 100,000, 30,000, 10,000. Nine batches were assigned for each simulation: six batches with two aliquots each and three batches with a single aliquot each. Two aliquots from each participant were assigned to the same batch, one aliquot from each of two randomly chosen participants were assigned to a batch together, and the remaining three aliquots (each from a separate participant) were assigned to the singleton batches. A separate random choice of participants for the mixed batch was made for each simulation.

In the scenario with no variation, each well is positive with probability 1 − exp (−*n* exp(*v_i_*)), where *n* is the number of input cells and *v_i_* is the log of infection frequency for participant *i*. In each scenario with variation, two different formulations were tested. *Normal formulation*: Random effects *a_ij_* and *b_kl_* were sampled from normal distributions (3), and each well is positive with probability 1 − exp (−*n* exp(*v_i_* + *a_ij_* + *b_kl_*)). *Gamma formulation*: Batch random effect *b_kl_* was sampled as in the normal formulation, and each well is positive with probability 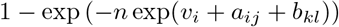, representing the probability that a negative binomial random variable is positive when it has mean *n* exp (*v_i_* + *b_kl_*) and dispersion parameter 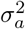.

For multi-lab simulations including both fresh and cryopreserved samples, the full experiment as specified in S3 Table was simulated, using ensemble posterior median parameter estimates reported in S8 Table (“All four labs, using JHU day 21 readout”). Random effects and the probability that each well is positive followed the model described above (“Statistical model”). In addition, to compute the probability that the method may overestimate *σ_a_, σ_b_*, or *σ_c_* despite the true parameter value equaling zero, we repeated the simulations three times, each time replacing one of these three parameters with zero. We then reported the average probability of overestimation among these simulations.

2000 simulations of each specification were performed, except for the analysis considering true parameter values of zero, for which only 100 were performed. ML estimation (above, “Maximum likelihood estimation”) was used for data resulting from single-lab simulations, and MCMC estimation (above, “Markov-chain Monte Carlo estimation”) was used for both single- and multi-lab simulations.

#### Characterizing accuracy of assays

Simulations were used to illustrate the typical accuracy of each lab’s assay in the absence of batch effects (assuming that these could be rendered irrelevant by assaying specimens from the same person — such as baseline and post-treatment — in the same batch). For each of 1000 samples from the batch effect-free ensemble posterior (see above, “Markov-chain Monte Carlo estimation”) and each assumed true IUPM, 1000 simulations of well-by-well output were performed for each assay. For each simulation, aliquot-level random effects *a_ij_* were sampled from 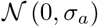, and lab-level random effects *c_il_* were either sampled from 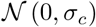, sampled from 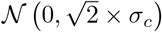, or set to zero (choice of distribution described below). In each simulation, the maximum likelihood estimate of IUPM was compared to the true IUPM (scaled by lab-specific factor *β_l_*) to obtain the log_10_-error. The median of absolute log_10_-error was then recorded for each of the 1000 samples from the ensemble posterior. For each lab’s assay, we then report the median, 2.5%ile, and 97.5%ile of these median absolute errors across the posterior sample.

We ran two versions of these simulations. For the first version, we supposed that each assay has equal claim to biological truth and that lab-based random effects *c_il_* are discrepancies from a consensus standard. These random effects were drawn from 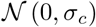, where *σ_c_* is the estimated value drawn from the ensemble posterior. For the second version, we chose the JHU assay (index *l* = 3) to represent the standard against which other assays are measured. Under this assumption, the differences *c_il_* − *c*_*i*,3_ are lab *l*’s discrepancies from the standard. To simulate data in which the *c_il_* themselves are the discrepancies, we sampled these random effects from 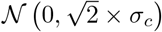 for all *l* ≠ 3. All *c*_*i*,3_ were set to zero.

The same simulations were also used to compare accuracy between each pair of assays. For each assumed true IUPM and ensemble posterior sample, the difference in median absolute log_10_-error between two assays was recorded for the 1000 matched simulations. For each pair of assays, we report the median, 2.5%ile, and 97.5%ile of these differences across the posterior sample.

At JHU and SR, the number of replicate wells with one million cells apiece varied with the number of rCD4s purified from each aliquot. First, to mimic typical aliquots, we simulated the JHU assay with 26 replicates and the SR assay with 12 replicates. Second, to consider the situation where JHU and SR used a number of cells similar to those used by U. Pitt. and UCSD, we simulated both assays with 8 replicates.

#### Simulation of clinical trial

Latency-reducing treatment was simulated to illustrate the power and accuracy of QVOA for measuring changes in IUPM. Pretreatment log_10_ IUPM from each participant was assumed to follow a normal distribution with mean −0.013 and standard deviation 0.675, consistent with the cohort studied in [10]. Within-person change in log10 IUPM from pre- to post-treatment was normally distributed with mean δ and standard deviation *σ_δ_*. In control participants, “pre” and “post” IUPMs were equal. Both JHU and UCSD assay protocols were simulated, using ensemble posterior median parameter estimates reported in S8 Table (“All four labs, using JHU day 21 readout”). For the JHU assay, we simulated 50 million-cell wells for fresh samples, 26 million-cell wells for frozen samples, and two wells of each other size (five-fold dilution from 200,000 to 320). The UCSD assay was simulated according to the configuration in S3 Table.

Both batched and unbatched analyses were studied. In batched analysis, all samples were treated as cryopreserved, and only aliquot- and lab-level excess variation were included (all batch effects *b_kl_* =0). In unbatched analysis, all samples were treated as fresh, and all three levels of excess variation were included. In both cases, the same lab-level random effect *c_il_* was assumed to hold for both samples from each participant. For each sample, the maximum likelihood IUPM was estimated, with negative assay results replaced with a value equal to one-half of the minimum possible positive assay value. A before-after change in log10 IUPM was calculated for each individual from these IUPM estimates.

Two treatment effects were studied: a 10-fold median reduction in IUPM (strong effect), but with substantial person-to-person variation (*δ* = −1, *σ_δ_* = 0.5), and a weak effect with lower person-to-person variation (*δ* = −log_10_(3) ≈ −0.48, *σ_δ_* = 0.16). We intended the strong effect to represent a treatment with clinical potential, and we simulated an early, discovery-oriented study with 6 treated participants and no controls, analyzed by a paired t-test. We intended the weak effect to represent a treatment with minor improvement over those currently reported [18], of interest for providing information about the biology of HIV or as a possible component of future combination treatments. We assumed greater concern in this situation about background change over time and assay drift, so we simulated studies with 12 treated and 12 control individuals, with change in log10 IUPM analyzed by unpaired t-test. We also evaluated an alternative approach to analysis of the simulated data. This approach used maximum likelihood applied to all the well-level assay results to estimate a model with fixed effects for treatment and each person’s baseline IUPM, along with a random effect for within-person change. (For the weak effect scenario, different random effect variances were allowed for treated versus control participants.) 1000 simulations were analyzed for each scenario, for 8000 simulations in all (JHU/UCSD assay protocols, batched/unbatched analyses, weak/strong treatment effect). Power was calculated as the fraction of simulations for which *p* < 0.05 in the t-test or maximum likelihood analysis.

## Results

### Simulation validates MCMC method

We first applied MCMC estimates to single-lab frozen panel simulations using the model with both aliquot- and batch-level random effects. We simulated assays encompassing 26 million cells from each of five participants, divided into three aliquots, and assayed as six replicates per aliquot in ~3-fold dilutions spanning 1 million to 10,000 cells. In the scenario with large variation, this method accurately estimated *σ_a_*. For example, applying a 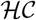(0, 2) prior to a simulation with normally distributed random effects, the estimate had median error +0.03 and 97.4% coverage of the 95% CI. Batch-level variation σ⅛ was subject to larger overestimates and overly conservative CIs, with median error +0.09 and 99.3% coverage. To control overestimates, which were worse in simulations with less variation, we relied on an ensemble model when studying multi-lab simulations and actual data. This approach allows for the possibility that one or more random effect sizes are zero, giving it prior probability of 0.5. Analysis by maximum likelihood gave poorer results, with substantial bias toward random effect variances of zero and poor coverage probabilities. Results for both methods were similar with the alternative gamma distribution assumption for the *a_ij_*.

In multi-lab simulations mimicking the full experimental design, we found that MCMC using a 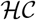(0,1) prior on random effect standard deviations adequately estimated all fixed effects and the between-lab random effect, with median biases (on the natural log scale) between −0.030 and +0.028 and 95% CI coverage between 93.2% and 95.0% (Table 2). While the combined effect of aliquot- and batch-level variation was estimated well (median bias −0.085, 95% CI coverage 96.6%), individual estimates performed less well: Aliquot-level variation was overestimated (median bias +0.099), batch-level variation was substantially underestimated (median bias −0.351), and both had low coverage. Inability to disentangle variation at both levels likely follows from the fact that the experiment was limited to batches of only one or two aliquots each, as occurred in the actual study. Experimental techniques that allow study of larger batches may help to reduce this error. Overall, the 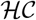(0,1) prior slightly outperformed the other three priors tested (S5 Table, S6 Table, S7 Table), and so it was chosen for all analysis of experimental data.

**Table 2.** Performance of MCMC estimation, using the ensemble model and 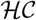(0,1) prior, in simulation of multi-lab experimental design in S3 Table (total of 194 million to 289 million cells from each of five participants, distributed among four labs, encompassing 474 to 569 wells per participant).

| Parameter | Median bias | Median absolute error | Interquartile coverage | 95% CI coverage |
| --- | --- | --- | --- | --- |
| Total variation<br>($\sqrt{\sigma_a^2 + \sigma_b^2 + \sigma_c^2}$) | +0.019 | 0.093 | 50.3% | 95.6% |
| Aliquot & batch variation<br>($\sqrt{\sigma_a^2 + \sigma_b^2}$) | -0.055 | 0.134 | 54.9% | 96.6% |
| Aliquot & lab variation<br>($\sqrt{\sigma_a^2 + \sigma_c^2}$) | +0.137 | 0.141 | 28.9% | 78.7% |
| Batch & lab variation<br>($\sqrt{\sigma_b^2 + \sigma_c^2}$) | -0.057 | 0.145 | 49.1% | 95.3% |
| $\sigma_a$ | +0.099 | 0.188 | 33.2% | 88.4% |
| $\sigma_b$ | -0.466 | 0.466 | 30.5% | 89.5% |
| $\sigma_c$ | +0.021 | 0.262 | 47.7% | 93.2% |
| $\beta_s$ | -0.000 | 0.219 | 50.7% | 95.0% |
| $\beta_2$ (UCSD) | +0.028 | 0.290 | 45.0% | 93.4% |
| $\beta_3$ (JHU) | +0.000 | 0.311 | 47.4% | 93.2% |
| $\beta_4$ (SR) | -0.030 | 0.326 | 49.2% | 94.0% |

To investigate whether MCMC might lead us to conclude falsely that variation is present at a given level, we simulated data for which one or more of the three sources of variation was removed. When all three sources were removed from simulations, MCMC rarely produced estimates surpassing the level of variation observed in the experimental data (posterior probability weight typically no more than 0.1%, bottom row of Table 3). When one or two sources of variation were included, MCMC often misidentified the source of variation, though misidentification as aliquot-level variation was less common and typically produced estimates smaller than the experimental estimate. Even when one source was misidentified as another, estimates of total variation coming from all three levels performed well.

**Table 3.**
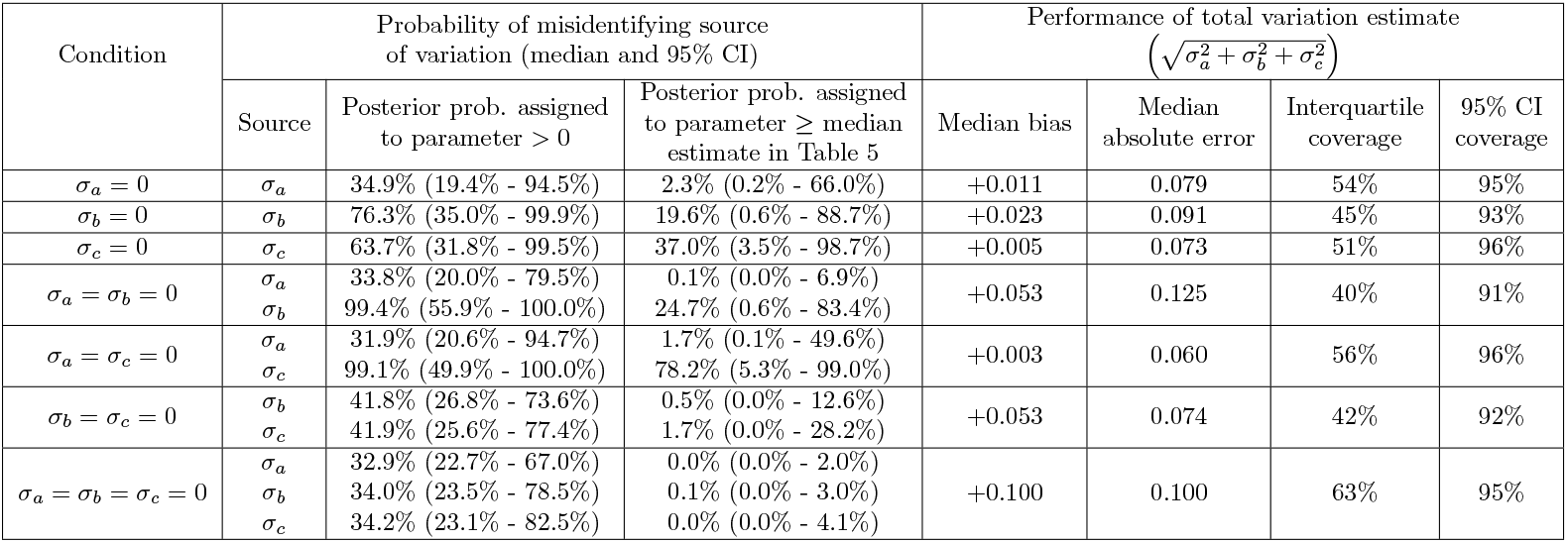
Performance of MCMC estimation assuming a true random effect parameter of zero, using 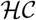(0,1) prior, in multi-lab simulation.

### MCMC method estimates systematic and random variation in QVOA

Although the goal of each lab’s assay is the same — to quantify infectious provirus infecting resting CD4^+^ T cells — infection levels reported by UCSD were consistently higher than those reported by the other three labs (Figure 2). We used the Bayesian model to estimate the systematic effect of assay characteristics and lab practices, measured as fold-change from U. Pitt. as reference (Table 4). When accounting for differing cell counts in each assay, as well as excess random variation at the aliquot, batch, and lab levels, we found that UCSD reported IUPMs averaging 9.2-fold higher than those reported by U. Pitt. (95% CI 3.8 – 24). For the other two labs, credible intervals for this systematic effect spanned 1. This result is not surprising, given methodological differences in each assay. U. Pitt., JHU, and SR each recorded a well as positive for viral outgrowth if levels of viral protein p24 measured by ELISA exceeded that of a threshold reference sample. UCSD, on the other hand, used viral RNA detection. While RNA detection is more sensitive than the p24 assay, it may have greater potential for false positive results. In fact, two wells of a single negative coded control aliquot were reported as positive (Figure 2).

**Fig 2.**
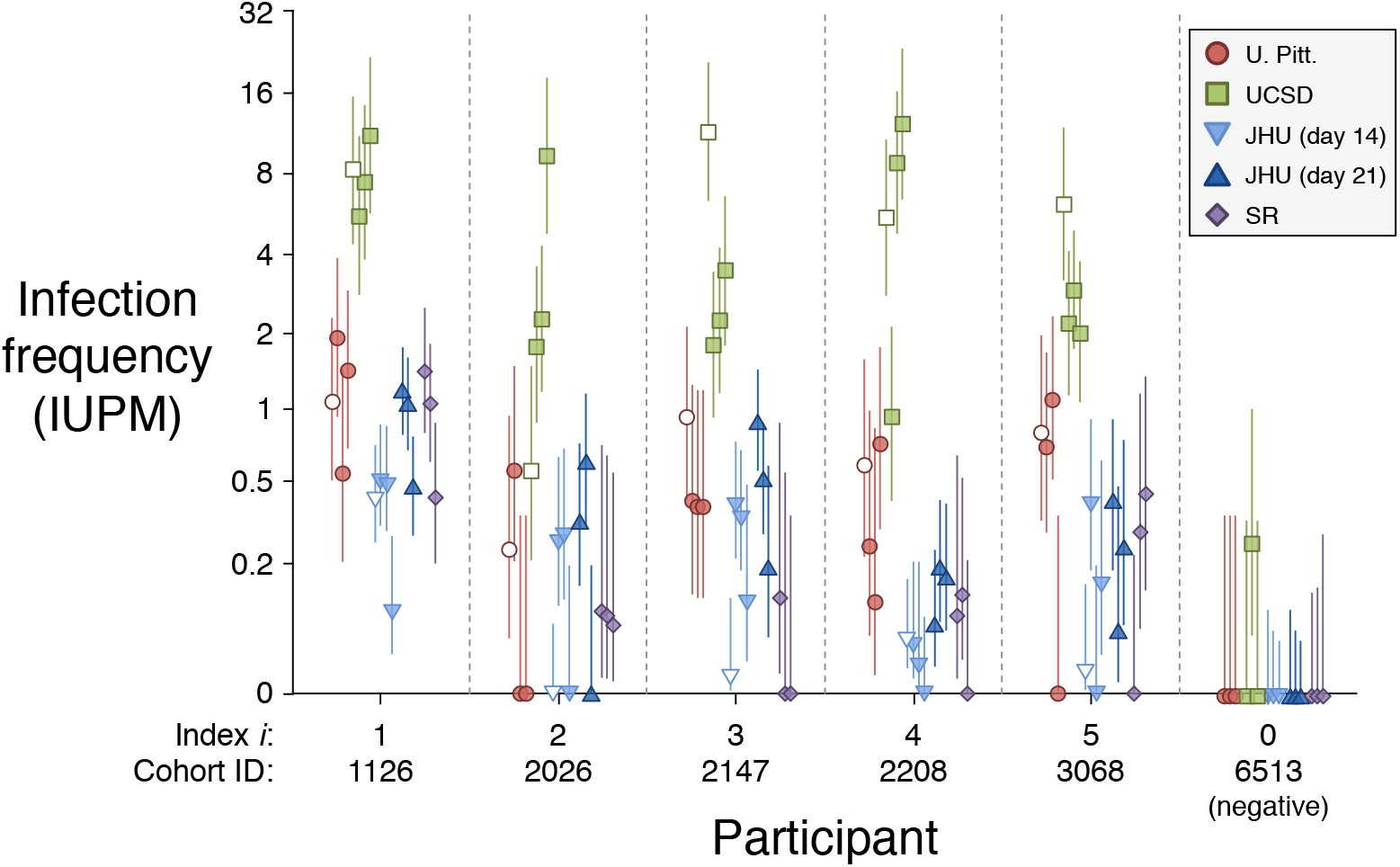
Maximum likelihood estimates and 95% CI of infection frequency for each aliquot. Cryopreserved aliquots are indicated by shaded symbols, fresh aliquots by open symbols. “Index *i*” is used in model output, and “Cohort ID” represents the identifier used in the SCOPE/OPTIONS cohort.

**Table 4.** Systematic lab effects, fold-change from U. Pitt. Posterior medians and 95% CIs shown.

| Lab | Ensemble model (accounting for excess variation) | Enforcing model without excess variation |
| --- | --- | --- |
| UCSD | 9.2 (3.8 - 24) | 7.4 (5.7 - 9.7) |
| JHU | 0.81 (0.30 - 2.4) | 0.75 (0.56 - 1.0) |
| SR | 0.39 (0.13 - 1.12) | 0.51 (0.35 - 0.75) |

Both the systematic effect and assay input cell counts determine the frequency of all-negative assay results. While SR’s experimental protocol was modeled on that of JHU (S2 Table), they reported more all-negative aliquots (4 of 15 versus 1 of 15 aliquots for JHU). This observation reflects the fact that SR generally recovered fewer resting CD4s from each aliquot than JHU did for input into QVOA (average of 12.7 versus 26.8 million cells per aliquot, S3 Table), and it might also reflect a systematically lower IUPM scale (half that of JHU, although credible intervals overlap).

In addition to this systematic lab effect, we determined that random variation in excess of the baseline Poisson-binomial model was likely (posterior probability 85%) at all three levels — between-aliquot, between-batch, and between-lab. Excess variation in at least one level was a near-certainty (posterior probability = 1 − 10^−22^). Table 5 summarizes estimates of excess variation at each level. As study design was limited to batches of only one or two aliquots, it is not easy to disentangle variation at the aliquot and batch levels; estimates are inversely correlated with one another (slope −0.69, S1 Fig). Combining variation at both of these levels, we estimate that two aliquots, studied in two different batches at the same lab, are expected to vary 2.0-fold in excess of Poisson variation (95% CI 1.6 – 2.7). Two aliquots, studied at two different labs, are expected to vary 2.3-fold in excess of Poisson variation (95% CI 1.8 – 3.5), an estimate obtained by combining variation at all three levels.

**Table 5.** Estimated excess variation in QVOA.

| Level | Posterior probability of excess variation |  | Estimated variation, fold-change (posterior median and 95% CI) |
| --- | --- | --- | --- |
| Between aliquot | 98.0% |  | 1.5 (1.1 – 2.1) |
| Between batch | 90.4% |  | 1.8 (1.0 – 2.4) |
| Between lab | 96.3% |  | 1.5 (1.0 – 2.5) |
| Between aliquot & batch, combined | $1 - 10^{-18}$ (var. at either level) | | 2.0 (1.6 – 2.7) |
| Total excess variation between aliquots in two different labs | At least one level | $1 - 10^{-22}$ | 2.3 (1.8 – 3.5) |
|  | At least two levels | 99.48% |  |
|  | All three levels | 85% |  |

### Cryopreservation did not have a major effect on QVOA outcomes

When accounting for excess variation, all credible intervals estimated for the effect of cryopreservation spanned 1-fold change, or the absence of an effect (Table 6). When using results from all three labs that tested fresh and frozen panel aliquots (SR did not test fresh PBMC samples) to estimate a single effect size, we estimated between 0.56- and 1.97-fold change in infection frequency compared to fresh aliquots. In the context of the large (> 100-fold) reductions sought by latency-reducing therapies, this effect is not major. Credible intervals were wider when the effect was estimated for each lab separately, and the interval was particularly wide for JHU (25-fold difference between top and bottom of interval), owing to the larger number of all-negative aliquots at this lab.

**Table 6.** Effect of cryopreservation, fold-change (posterior median and 95% CI).

| Lab(s) | Ensemble model (accounting for excess variation) | Enforcing model without excess variation |
| --- | --- | --- |
| All four, using JHU day 14 readout | 1.04 (0.56 – 1.97) | 1.0 (0.78 – 1.26) |
| U. Pitt. | 0.61 (0.18 – 1.65) | 0.71 (0.42 – 1.18) |
| UCSD | 0.79 (0.27 – 2.17) | 0.75 (0.56 – 1.05) |
| JHU (day 14 readout) | 2.89 (0.62 – 15.7) | 2.09 (1.30 – 3.44) |

### Precision within and across labs

Running MCMC analysis on each lab separately suggested that each lab’s assay offered a similar level of precision. The median estimates for aliquot- and batch-level variation for each lab fell within the 95% credible interval of the joint estimates in Table 5, and none were more than 21% away from the corresponding median estimate (S8 Table, S2 Fig). Precision of early readout (day 14) from JHU assays was, however, estimated to be lower than that of the other assays: There was combined 3.5-fold variation at both levels (95% CI 2.0-to 10.7-fold), which is 69% higher (95% CI 10% smaller to 458% higher) than the joint lab estimate that included data from the later JHU readout. This difference in precision may reflect the fact that allowing more time for exponential growth leads to a stronger p24 signal and clearer distinctions between positive and negative wells. S9 Table provides full joint posteriors for each separate analysis.

Neglecting the strong evidence supporting excess random variation at multiple levels (Table 5) can generate misleading interpretations about precision of lab procedures. To demonstrate the relevance of accounting for excess random variation, we recomputed all estimates in a model excluding this excess variation (Tables 4 and 6, right column). While median estimates did not change greatly, credible intervals shrank by 2-to 10-fold for each parameter. Neglecting this excess variation therefore overstates certainty in parameter estimates. One particular effect of this error in our experiment would be to conclude, rather strongly (*p* < 0.001), that cryopreservation increases observed infection frequencies in the JHU lab.

Paying close attention to the sources of random and systematic variation can help in choosing assays for and optimizing design of clinical trials for latency-reducing therapies. In the next two sections, we demonstrate how simulations based on the parameter estimates described above may guide this effort.

### Simulations based on experimental outcomes can be used to compare assay accuracy

While each lab’s assay aims to measure the replication-competent HIV latent reservoir, protocols differ among them: U. Pitt., JHU, and SR use a p24 antigen test to detect viral outgrowth, while UCSD uses an RNA PCR test; U. Pitt., JHU, and SR use PHA and gamma-irradiated PBMCs to stimulate resting cells, while UCSD uses antibody to CD3/CD28 bound to the culture plate (S2 Table). Additionally, target cells added to propagate virus differs among labs. These protocol differences may explain both systematic and random variation between labs (Tables 4, 5). We may think of these labs as measuring different aspects of latency, each with a valid claim to being a meaningful measure, with experimental and biological motivations for specific protocol choices.

In the absence of an external standard defining latent reservoir size, we can nonetheless address how sensitivity of an assay affects its accuracy. Specifically, assays that use more input cells overall or have systematically high IUPMs (high fold-change in Table 4) will have improved sensitivity. By drawing from the joint posterior distribution (S9 Table) and simulating data for each draw, we investigated how sensitivity relates to the accuracy of measuring small reservoirs (see Methods).

For typical infection frequencies (1 IUPM on the U. Pitt. scale, higher or lower for the other assays according to systematic effect, Table 4), all four assays have nearly identical accuracy, as measured by median absolute error from a consensus standard (Figure 3). As infection frequency declines from this level, error increases for all four assays, most sharply for SR, but gradually for UCSD and JHU. In the case of SR simulated at 0.1 IUPM on the U. Pitt. scale (median of 0.039 IUPM on the SR scale, Table 4), for a majority of parameter draws, a majority of simulated assay outcomes are all-negative, resulting in infinite error on the log scale. In the case of UCSD and JHU, improved accuracy has costs: The more sensitive RNA-based assay used by UCSD may have a higher false positive rate (2 of 18 control wells with one million cells were reported as positive, compared to none of the 193 control wells with one million cells among the other three assays, S4 Table), and JHU exhausted a larger sample (26.5 million cells simulated, versus 8.6 to 12.5 million for the other assays). If the JHU assay is simulated with fewer cells, bringing it in line with the other assays, its accuracy profile lies between that of U. Pitt and SR (S3 Fig); overlapping credible intervals in accuracy reflect overlapping credible intervals in systematic lab effects.

**Fig 3.**
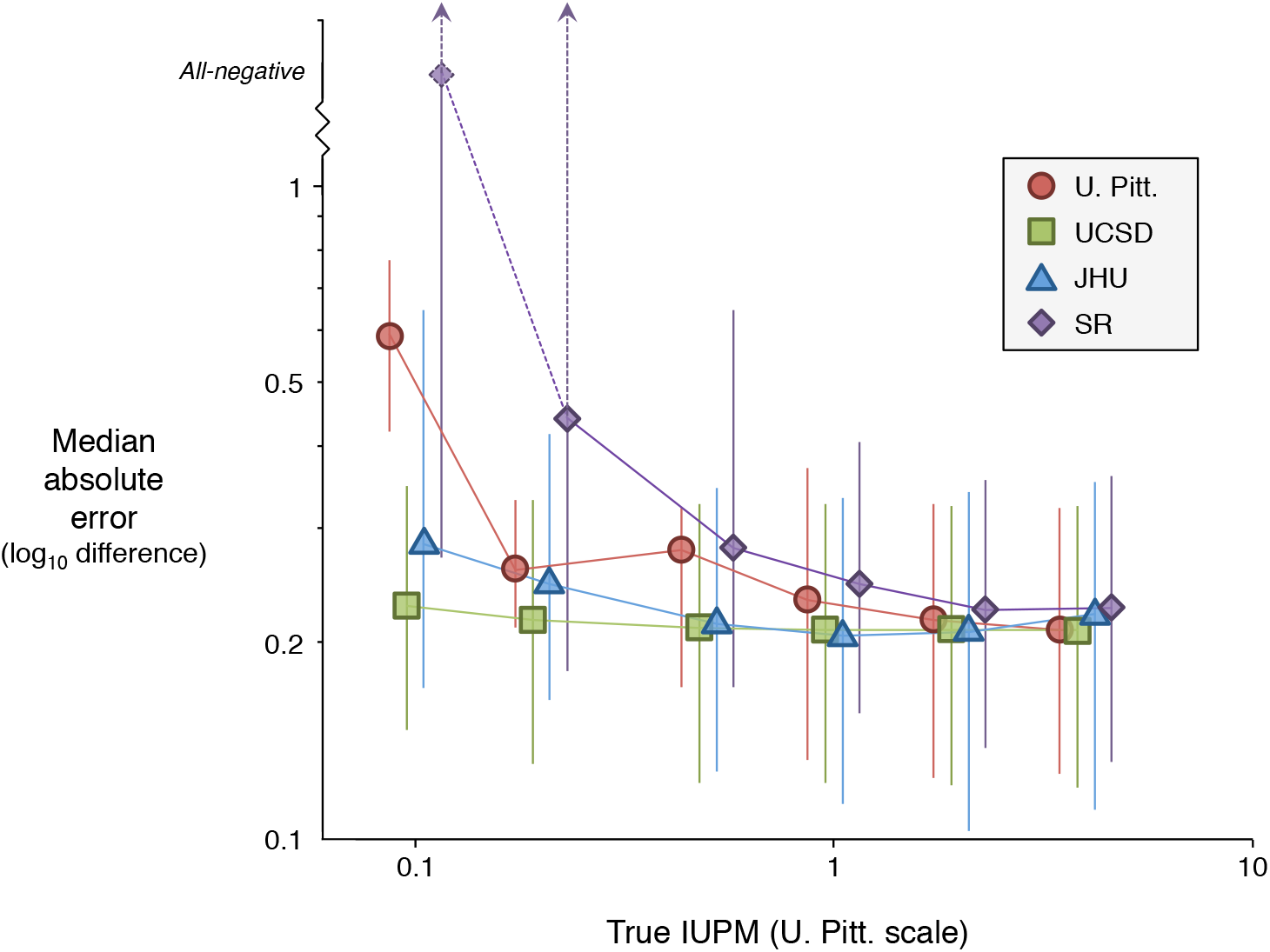
Accuracy of assays used in the experimental study. Each assay is measured against a consensus standard, appropriately scaled by *β_l_* for that assay. “All-negative” represents infinite error on the log scale, which occurs when the maximum likelihood estimate of IUPM is zero.

If false positives did in fact occur in HIV^+^ samples studied by the UCSD RNA-based assay, it would be reflected in our estimates as both improved sensitivity for UCSD (higher *β*_2_) and reduced correlation between labs (higher *σ_c_*). We used simulations to investigate the possible tradeoff between sensitivity and specificity, treating the JHU assay as a gold standard (not subject to lab-based random effect) and subjecting UCSD to the total between-lab variation (applying a random effect with standard deviation 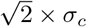 to each participant). If an equal number of cells is made available to both assays, then the extra sensitivity of RNA readout outweighs the cost of extra variability for particularly small reservoirs (IUPM of 0.2 or less on the U. Pitt, scale, Figure 4). Uncertainty in estimates of between-lab variation, UCSD systematic effect, and JHU systematic effect, however, make definitive comparisons difficult; credible intervals for the UCSD/JHU accuracy difference overlap zero for all IUPMs simulated. If the JHU assay instead uses roughly three times as many cells as UCSD, as in the experimental study, then it does not suffer the same disadvantage at the low IUPMs simulated and enjoys a somewhat larger advantage at higher IUPMs. The accuracy of any two assays may be compared head-to-head by the same simulation method, judging their performance against a consensus standard (applying *σ_c_* equally to both assays, S10 Table and S11 Table) or against a chosen standard assay (applying 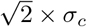 to other assays, S12 Table and S13 Table).

**Fig 4.**
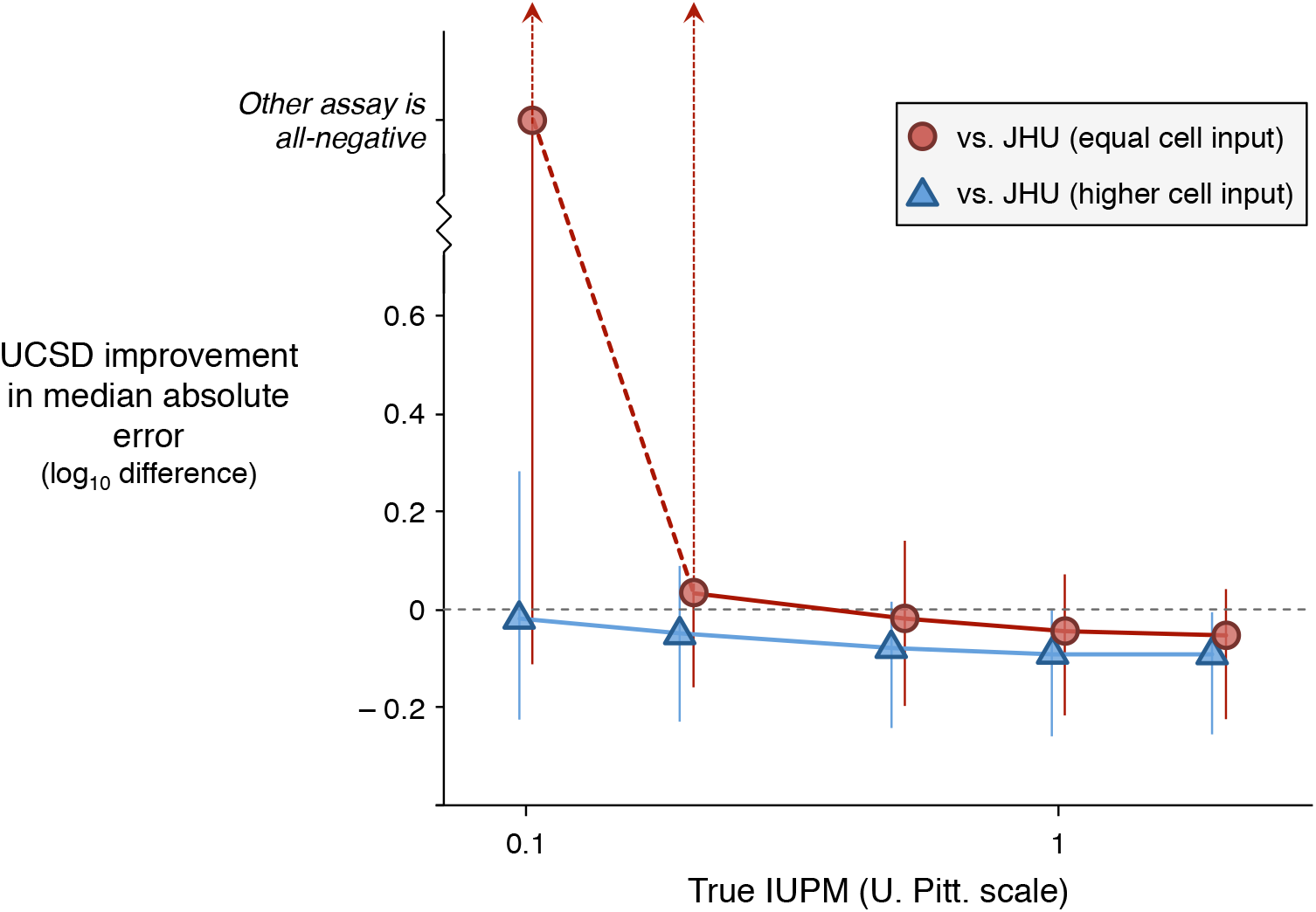
Difference in accuracy between UCSD and JHU assays, assuming that the JHU assay is a gold standard (not subject to lab-based random effect). Batch variation-free ensemble estimates of parameters were used in simulations. Median estimate and 95% credible intervals shown for 0.1, 0.2, 0.5, 1, and 2 IUPM on the U. Pitt, scale. All values plotted are also provided in S12 Table and S13 Table.

### Batched study design can improve power to detect latency reduction

Cryopreserving specimens allows samples taken at different points in time to be thawed and analyzed together in the same batch. This strategy eliminates the estimated 1.8-fold batch variation that would otherwise affect longitudinal comparisons in clinical trials, potentially boosting power to detect reduction in latency. To investigate this possibility, we simulated and analyzed data for two hypothetical latency-reducing therapies with strong effect (10-fold reduction in latency) and weak effect (3-fold reduction), as described in Methods. For each therapy, we simulated both a p24 assay based on JHU protocol, which uses all available cells from a participant, and an RNA assay based on UCSD protocol, which uses a fixed number of cells regardless of availability. Consistent with JHU assay cell counts (S3 Table), we supposed that roughly half as many cells would be recovered from cryopreservation as would be available fresh.

For the weaker therapy, batching substantially improved power to detect and accuracy to measure latency reduction following treatment (Table 7). Improvement was greater for the UCSD protocol, with power increasing from 58.6% to 82.6% and median absolute error declining from 0.14 log_10_ to 0.10 log_10_. Although freezing-thawing sacrificed half of the resting cells available to the JHU protocol, the benefits of batching overcame this deficit, leading to a 15 percentage-point increase in study power.

**Table 7.**
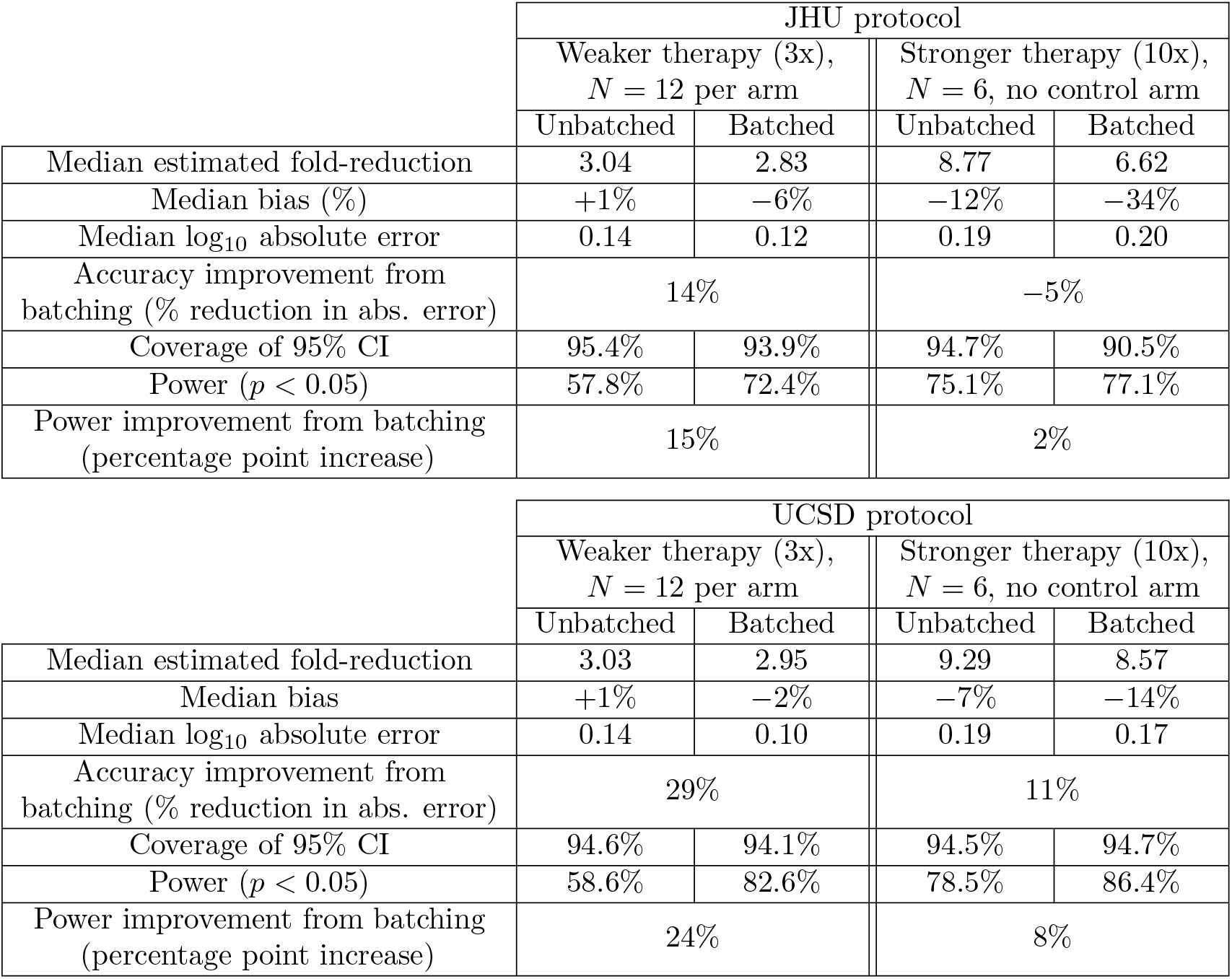
Simulated results of latency reduction trials, using a t-test to compare pre- and post-treatment data

For the stronger therapy studied in a smaller cohort, batching provided a modest benefit to the UCSD protocol, but reduced cell availability in the JHU protocol eliminated this benefit completely. Moreover, both protocols underestimated the effect of treatment (−7% to −34% bias (Table 7). This outcome resulted from our use of an imputed IUPM in the case of negative assay results, a common practice in the field [19]. Strong therapy that reduces latency 10-fold can often result in true IUPMs well below the imputed value. In the batched JHU protocol, 35% of samples returned a negative assay, leading to the largest underestimate of treatment effect. To provide an alternative to the t-test that avoided imputation, a maximum likelihood estimate of effect size was also computed. When this method was used, batching improved both power and accuracy in all cases, but the resulting confidence intervals had poorer coverage (S14 Table).

### Maximum likelihood estimation underperformed MCMC

None of the three software packages that we tried produced fully satisfactory results when applied to the experimental data. The best log-likelihood obtained was from a model in Stata that excluded between-lab variation. When this variation was included, the log-likelihood closely approached that of the best model before ending with an error. In SAS, the model fitting completed for essentially the same model, with *σ_c_* = 0, but with most parameters having standard errors equal to zero. In R, the model converged to substantially different parameter values, with a much worse log-likelihood.

Aside from the lack of between-lab variation, which may reflect the bias toward zero seen in single-lab simulations (see “Simulation validates MCMC method,“ above), many results by maximum likelihood were similar to the MCMC results given above. The cryopreservation effect and systematic lab effects were all within 10% of MCMC posterior medians. The typical between-aliquot excess variation was estimated to be 1.61-fold (vs 1.50-fold by MCMC), with a p-value of 0.0003. The typical between-batch excess variation was estimated to be 1.43-fold (vs 1.77-fold by MCMC), with *p* = 0.098. A likelihood ratio test for at least one source of excess variation had *p* =1.4 × 10^−13^.

## Discussion

Rigorous evaluation of the performance of QVOAs is challenging due to their high cost and the presence of unavoidable background variation. By applying detailed statistical analysis to 75 split PBMC samples from 5 ART-suppressed HIV^+^donors, we found strong evidence for additional variation beyond the unavoidable background, even within a single batch at a single lab. When a donor’s true IUPM is 1 or more (on the U. Pitt. scale; Table 4), we estimated that assay results typically differ from the true IUPM by a factor of 1.6 to 1.9 up or down (Figure 3, typical log_10_ difference of 0.20 to 0.28). At lower true IUPMs, some assays had much larger median errors, while others avoided major increases by use of a large number of input cells (JHU assay) or by use of a readout that was more often positive (UCSD assay).

We found that the four assays appeared to differ from each other both systematically (in scale; see Table 4) and randomly (between-lab variation in Table 5). These differences may reflect the differing subsets of infectious units (IU) that the assays detect. Recent work suggests that QVOAs detect only a fraction of the IU present in a sample [11]. The assays’ different stimulation and detection procedures likely cause each to measure a different subset of IU. Different inducers of T cell activation have variable effectiveness. In one study, the use of antibodies to CD3 and CD28 was more effective than other methods [20]. For the detection step, HIV RNA by PCR is more sensitive than digital ELISA assays for p24 antigen (e.g., Simoa [21]), which in turn is more sensitive than the standard ELISA assay for p24 antigen. Details of CD4 cell separation and aspects of cell co-culture details like cell type and medium could also create variability between labs. Measuring a larger subset of IU is desirable, because sampling variation is relatively smaller when counts are higher, reducing relative error. A caveat, however, is that the fraction of the total latent reservoir within a smaller subset could, at least in principle, be more stable within and between individuals than that within a larger subset. We could not assess this possibility, because we had no gold standard measurement of the entire reservoir. In addition, higher readings that are due to false positive wells will not improve measurement. The UCSD assay, which used an in-house RT-PCR assay to detect HIV RNA rather than p24 antigen in culture supernatants, had two false positive wells among 126 wells evaluated for an HIV-uninfected donor. These were contiguous wells with one million cells each within a single one of three split samples. This may have been due to contamination during culture supernatant testing by RT-PCR, and in any case would not be a high enough rate to account for much of the UCSD scale factor.

The existence of substantial measurement error in QVOA has implications for the design of clinical studies that use QVOA to evaluate eradication strategies. Classical sample size calculations depend on the size of the latency reduction effect and how much it varies from person to person (*δ* and *σ_δ_* in the simulated clinical trials described above). Measurement error increases the effective person-to-person variation, which increases the sample size needed for any given level of desired precision. We do not, however, recommend assuming that measurement error is the *only* source of person-to-person variation in the observed effect. Interventions may well have truly differing effects for different people, and this variation may be as large or larger than the variation due to measurement error. Unfortunately, the variation in intervention effects may be difficult to accurately anticipate at the planning stage of a study [22].

Another implication is the need to use designs that will not be biased by regression-to-the-mean phenomena. For example, it might seem reasonable to exclude from an intervention study potential participants who have QVOA results that are all negative (measured IUPM = 0) at baseline, because they cannot go any lower. This exclusion, however, will tend to cause the remaining participants to average lower on repeat testing than at baseline, because the remaining subset has been depleted of some downward measurement errors by the exclusion, while the repeat testing will include an unbiased array of measurement errors. Therefore, a randomized parallel design should be used if such a baseline exclusion is applied; treated and control arms will then have comparable regression to the mean, resulting in a fair assessment of the intervention effect.

We found evidence that controlled-rate freezing and liquid nitrogen storage and shipping of PBMC did not cause substantial differences in QVOA results compared to use of fresh cells. This is encouraging because of the practical difficulties in using fresh cells and because storage enables assaying multiple specimens from the same person in the same batch, eliminating batch variation from estimates of within-person changes. It also permits clinical trials to be more readily conducted in trial sites without local laboratories qualified to perform reservoir assays. Our simulations showed the potential for gains in power and precision from such batching. There are, however, some important caveats regarding storage. We had only 15 results on fresh cells, with no split samples evaluated at the same lab. Despite this, our CI’s were narrow enough to provide good evidence against any systematic change due to storage of 2-fold or more (Table 6). In addition, the lack of split fresh samples precluded effective evaluation of the effect of storage on measurement variability. We also did not evaluate the effect of storage duration. The SR assay was performed later, after longer storage, than the other assays, and it also tended to have the lowest IUPMs. Using similar protocols to the JHU assay, it also obtained lower numbers of input cells, and JHU obtained lower input from stored samples than from the fresh samples collected at the same time. Consequently, until additonal studies on impact of duration of storage on QVOA IUPMs are conducted, we recommend that serial PBMC samples from participants in intervention studies be frozen and batch tested as soon a practical following collection.

Our statistical methods and results indicate potential value for additional statistical research on how to estimate and analyze QVOA data (and data from similar limiting dilution assays). We found a 98% posterior probability of extra-Poisson variation between split samples in the same batch at the same lab (Table 5). This suggests that there could also be extra-Poisson variability between wells within a single assay run, which would violate the single-hit Poisson dynamics that are typically assumed when calculating IUPM. A modified approach might be warranted in which the usual Poisson formulas are replaced with formulas based on the negative binomial distribution, which allows for overdispersion. A reasonable choice of overdispersion parameter would be our posterior median estimated 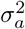 of about 0.16, or it could be estimated as an auxiliary parameter of the calculation. Alternatively, the usual maximum likelihood approach could be replaced with a Bayesian posterior median. For analysis of the full 75 QVOA results in our study, we found that Bayesian MCMC estimation outperformed maximum likelihood. Although our study differs from clinical studies of interventions to reduce the latent reservoir, there may be potential for MCMC estimation to improve the analysis of clinical studies, too.

## Conclusion

This study provides evidence that QVOA assays have extra variation, beyond what is theoretically inevitable, at three levels: between split samples even in the same batch, between batches run using the same assay, and between different assays. Results for stored frozen samples did not appear to differ systematically from those on fresh samples. We developed and validated methods for fitting detailed statistical models of assay variation. We are now using these methods to evaluate faster and cheaper alternatives to the classical QVOA assays evaluated here [23].

## Supporting information

**S1 Table Experimental setup, showing lab and batch assignment of each aliquot**.

**S2 Table.**
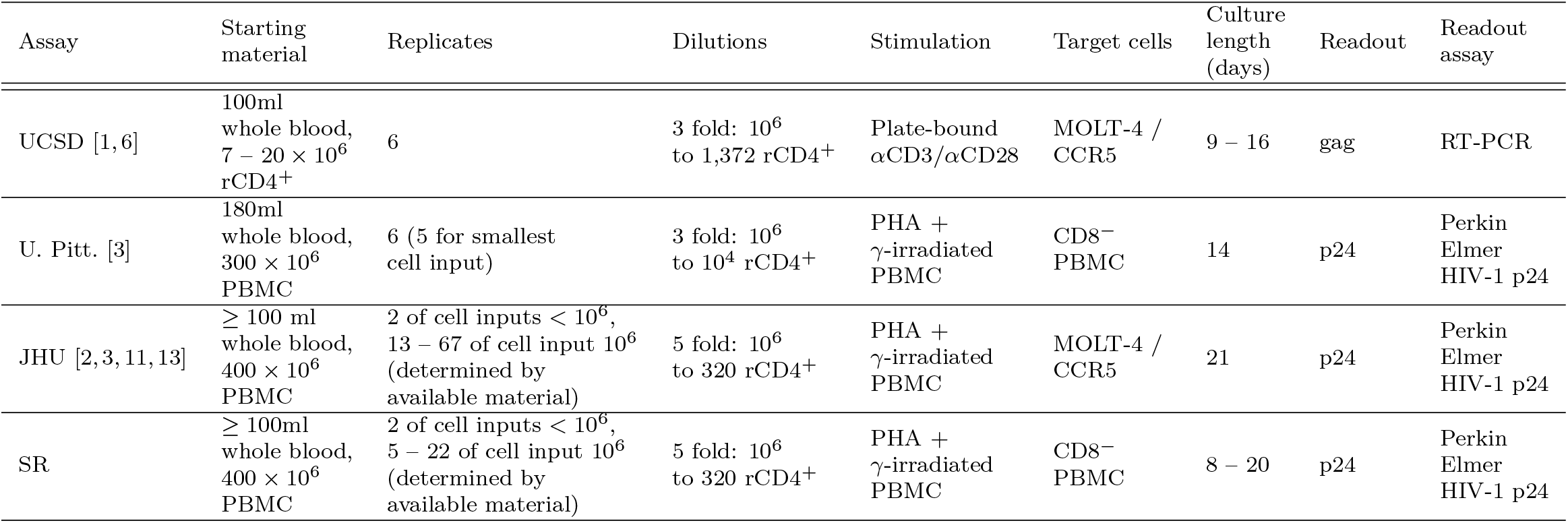
Laboratory protocols.

**S3 Table QVOA well configuration**.

**S4 Table Complete experimental dataset**.

**S5 Table.** Table Performance of MCMC estimation, using the ensemble model and 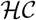(0,2) prior, in multi-lab simulation.

| Parameter | Median bias | Median absolute error | Interquartile coverage | 95% CI coverage |
| --- | --- | --- | --- | --- |
| Total variation<br>$\left(\sqrt{\sigma_a^2 + \sigma_b^2 + \sigma_c^2}\right)$ | +0.033 | 0.099 | 48.9% | 95.1% |
| Aliquot & batch variation<br>$\left(\sqrt{\sigma_a^2 + \sigma_b^2}\right)$ | −0.040 | 0.135 | 56.0% | 97.2% |
| Aliquot & lab variation<br>$\left(\sqrt{\sigma_a^2 + \sigma_c^2}\right)$ | +0.145 | 0.148 | 27.8% | 77.9% |
| Batch & lab variation<br>$\left(\sqrt{\sigma_b^2 + \sigma_c^2}\right)$ | −0.039 | 0.150 | 49.7% | 95.6% |
| $\sigma_a$ | +0.104 | 0.191 | 32.8% | 88.3% |
| $\sigma_b$ | −0.473 | 0.473 | 33.6% | 91.8% |
| $\sigma_c$ | +0.032 | 0.264 | 48.8% | 93.5% |
| $\beta_S$ | +0.002 | 0.219 | 50.8% | 95.0% |
| $\beta_2$ | +0.030 | 0.289 | 46.4% | 93.8% |
| $\beta_3$ | +0.001 | 0.311 | 48.6% | 94.0% |
| $\beta_4$ | −0.033 | 0.329 | 49.4% | 94.2% |

**S6 Table.**
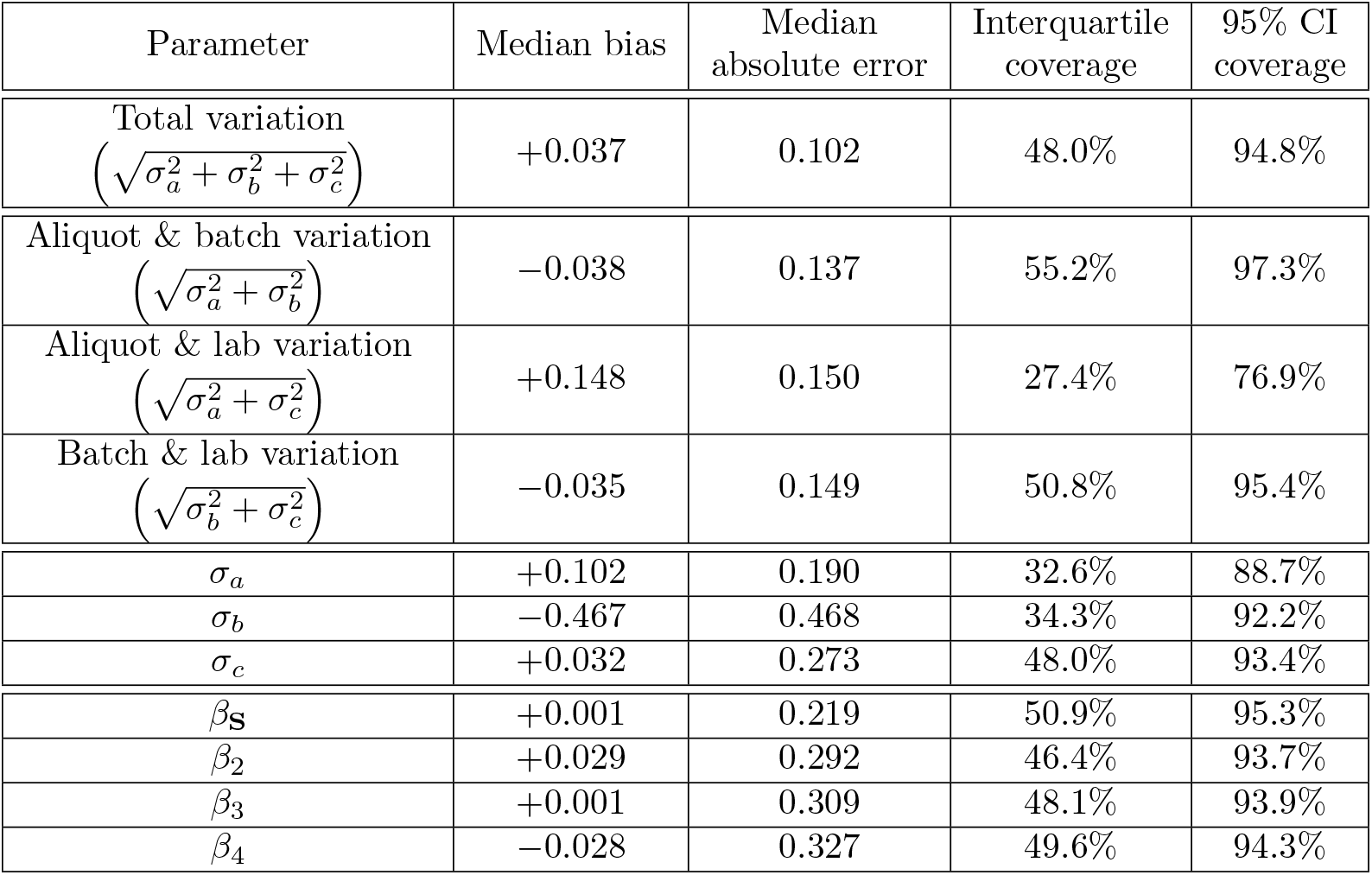
Table Performance of MCMC estimation, using the ensemble model and 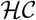(0,3) prior, in multi-lab simulation.

**S7 Table.** Table Performance of MCMC estimation, using the ensemble model and 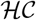(0,4) prior, in multi-lab simulation.

| Parameter | Median bias | Median absolute error | Interquartile coverage | 95% CI coverage |
| --- | --- | --- | --- | --- |
| Total variation<br>$\left(\sqrt{\sigma_a^2 + \sigma_b^2 + \sigma_c^2}\right)$ | +0.040 | 0.101 | 47.8% | 94.7% |
| Aliquot & batch variation<br>$\left(\sqrt{\sigma_a^2 + \sigma_b^2}\right)$ | −0.036 | 0.138 | 55.0% | 97.3% |
| Aliquot & lab variation<br>$\left(\sqrt{\sigma_a^2 + \sigma_c^2}\right)$ | +0.148 | 0.153 | 27.4% | 76.4% |
| Batch & lab variation<br>$\left(\sqrt{\sigma_b^2 + \sigma_c^2}\right)$ | −0.031 | 0.148 | 50.1% | 95.6% |
| $\sigma_a$ | +0.107 | 0.195 | 31.6% | 88.5% |
| $\sigma_b$ | −0.474 | 0.476 | 35.0% | 92.1% |
| $\sigma_c$ | +0.019 | 0.271 | 48.1% | 93.7% |
| $\beta_S$ | +0.000 | 0.221 | 51.1% | 95.5% |
| $\beta_2$ | +0.033 | 0.289 | 46.7% | 93.6% |
| $\beta_3$ | +0.002 | 0.311 | 48.3% | 94.1% |
| $\beta_4$ | −0.032 | 0.327 | 49.8% | 94.5% |

**S8 Table Summary of model fits**. Parameter distributions of the ensemble model are reported as weighted quantiles of 8000 posterior samples, and the number of samples in the discrete ensembles are shown (see Methods, “Markov-chain Monte Carlo estimation”). Analysis is shown for eight different subsets of the data: three subsets including multiple labs, and five subsets including a single lab each.

**S9 Table Full joint posteriors for model fits and membership in discrete ensemble posteriors**. Both discrete ensemble posteriors are provided for each subset of the data, each containing 1000 samples (see Methods, “Markov-chain Monte Carlo estimation”). Quantiles of both discrete ensembles are shown. Additionally, quantiles are shown for the single model estimating all three random effects.

**S10 Table.** Pairwise comparisons of assay accuracy for lower IUPMs, assuming that each assay has equal claim to biological truth (*σ_c_* applied to both assays). Batch variation-free ensemble estimates of parameters were used in simulations. Each entry shows median improvement of row versus column assay (median and 95% CI sampling from ensemble parameter distribution). IUPMs are shown on the U. Pitt. scale. Darker shading: Entire CI is above zero (green) or below zero (red). Lighter shading: Median estimate is above 0.05 (> 12% increase, green) or below −0.05 (> 11% decrease, red), but the CI crosses zero. “Infinite” difference in accuracy indicates that a majority of simulations of the disfavored assay have all-negative outcomes (maximum likelihood estimate of zero). Assay configurations match those in the experimental study, except “JHU (8M)” and “SR (8M),” which use fewer wells to match the cell input count of U. Pitt. and UCSD assays.

| Improvement in $\log_{10}$ error, IUPM = 0.1 | U. Pitt. | UCSD | JHU | SR | JHU (8M) | SR (8M) |
| --- | --- | --- | --- | --- | --- | --- |
| U. Pitt. | | -0.34<br>(-0.46 to -0.19) | -0.27<br>(-0.43 to 0.12) | $+\infty$<br>(-0.32 to $+\infty$) | 0.38<br>(-0.37 to $+\infty$) | $+\infty$<br>(-0.11 to $+\infty$) |
| UCSD | 0.34<br>(0.19 to 0.46) | | 0.06<br>(-0.04 to 0.40) | $+\infty$<br>(0.04 to $+\infty$) | 0.70<br>(0.00 to $+\infty$) | $+\infty$<br>(0.24 to $+\infty$) |
| JHU | 0.27<br>(-0.12 to 0.43) | -0.06<br>(-0.40 to 0.04) | | $+\infty$<br>(-0.01 to $+\infty$) | 0.59<br>(-0.02 to $+\infty$) | $+\infty$<br>(0.11 to $+\infty$) |
| SR | $-\infty$<br>($-\infty$ to 0.32) | $-\infty$<br>($-\infty$ to -0.04) | $-\infty$<br>($-\infty$ to 0.01) | | 0.00<br>($-\infty$ to $+\infty$) | 0.00<br>(0.00 to $+\infty$) |
| JHU (8M) | -0.38<br>($-\infty$ to 0.37) | -0.70<br>($-\infty$ to 0.00) | -0.59<br>($-\infty$ to 0.02) | 0.00<br>($-\infty$ to $+\infty$) | | 0.00<br>(-0.13 to $+\infty$) |
| SR (8M) | $-\infty$<br>($-\infty$ to 0.11) | $-\infty$<br>($-\infty$ to -0.24) | $-\infty$<br>($-\infty$ to -0.11) | 0.00<br>($-\infty$ to 0.00) | 0.00<br>($-\infty$ to 0.13) | |

| Improvement in $\log_{10}$ error, IUPM = 0.2 | U. Pitt. | UCSD | JHU | SR | JHU (8M) | SR (8M) |
| --- | --- | --- | --- | --- | --- | --- |
| U. Pitt. | | -0.05<br>(-0.09 to 0.00) | -0.02<br>(-0.08 to 0.11) | 0.18<br>(-0.06 to $+\infty$) | 0.06<br>(-0.08 to $+\infty$) | $+\infty$<br>(-0.04 to $+\infty$) |
| UCSD | 0.05<br>(0.00 to 0.09) | | 0.03<br>(-0.03 to 0.14) | 0.23<br>(-0.02 to $+\infty$) | 0.10<br>(-0.03 to $+\infty$) | $+\infty$<br>(0.01 to $+\infty$) |
| JHU | 0.02<br>(-0.11 to 0.08) | -0.03<br>(-0.14 to 0.03) | | 0.19<br>(-0.05 to $+\infty$) | 0.09<br>(-0.06 to $+\infty$) | $+\infty$<br>(-0.01 to $+\infty$) |
| SR | -0.18<br>($-\infty$ to 0.06) | -0.23<br>($-\infty$ to 0.02) | -0.19<br>($-\infty$ to 0.05) | | -0.08<br>($-\infty$ to $+\infty$) | 0.19<br>(-0.10 to $+\infty$) |
| JHU (8M) | -0.06<br>($-\infty$ to 0.08) | -0.10<br>($-\infty$ to 0.03) | -0.09<br>($-\infty$ to 0.06) | 0.08<br>($-\infty$ to $+\infty$) | | 0.46<br>(-0.15 to $+\infty$) |
| SR (8M) | $-\infty$<br>($-\infty$ to 0.04) | $-\infty$<br>($-\infty$ to -0.01) | $-\infty$<br>($-\infty$ to 0.01) | -0.19<br>($-\infty$ to 0.10) | -0.46<br>($-\infty$ to 0.15) | |

**S11 Table.** Table Pairwise comparisons of assay accuracy for higher IUPMs, assuming that each assay has equal claim to biological truth (*σ_c_* applied to both assays). Method and format match S10 Table.

| Improvement in $\log_{10}$ error, IUPM = 0.5 | U. Pitt. | UCSD | JHU | SR | JHU (8M) | SR (8M) |
| --- | --- | --- | --- | --- | --- | --- |
| U. Pitt. | | -0.06<br>(-0.09 to 0.00) | -0.05<br>(-0.10 to 0.02) | 0.01<br>(-0.08 to 0.34) | 0.00<br>(-0.08 to 0.15) | 0.04<br>(-0.09 to $+\infty$) |
| UCSD | 0.06<br>(0.00 to 0.09) | | 0.00<br>(-0.03 to 0.06) | 0.07<br>(-0.02 to 0.39) | 0.05<br>(-0.02 to 0.19) | 0.10<br>(-0.02 to $+\infty$) |
| JHU | 0.05<br>(-0.02 to 0.10) | 0.00<br>(-0.06 to 0.03) | | 0.06<br>(-0.03 to 0.40) | 0.05<br>(-0.03 to 0.19) | 0.09<br>(-0.03 to $+\infty$) |
| SR | -0.01<br>(-0.34 to 0.08) | -0.07<br>(-0.39 to 0.02) | -0.06<br>(-0.40 to 0.03) | | -0.02<br>(-0.34 to 0.12) | 0.04<br>(-0.16 to $+\infty$) |
| JHU (8M) | 0.00<br>(-0.15 to 0.08) | -0.05<br>(-0.19 to 0.02) | -0.05<br>(-0.19 to 0.03) | 0.02<br>(-0.12 to 0.34) | | 0.05<br>(-0.12 to $+\infty$) |
| SR (8M) | -0.04<br>($-\infty$ to 0.09) | -0.10<br>($-\infty$ to 0.02) | -0.09<br>($-\infty$ to 0.03) | -0.04<br>($-\infty$ to 0.16) | -0.05<br>($-\infty$ to 0.12) | |

| Improvement in $\log_{10}$ error, IUPM = 1 | U. Pitt. | UCSD | JHU | SR | JHU (8M) | SR (8M) |
| --- | --- | --- | --- | --- | --- | --- |
| U. Pitt. |  | -0.02<br>(-0.04 to 0.01) | -0.03<br>(-0.05 to 0.02) | 0.01<br>(-0.04 to 0.13) | 0.01<br>(-0.03 to 0.10) | 0.04<br>(-0.04 to 0.18) |
| UCSD | 0.02<br>(-0.01 to 0.04) |  | -0.01<br>(-0.03 to 0.03) | 0.03<br>(-0.02 to 0.16) | 0.03<br>(-0.01 to 0.12) | 0.06<br>(-0.02 to 0.20) |
| JHU | 0.03<br>(-0.02 to 0.05) | 0.01<br>(-0.03 to 0.03) |  | 0.04<br>(-0.02 to 0.16) | 0.04<br>(-0.02 to 0.12) | 0.06<br>(-0.02 to 0.20) |
| SR | -0.01<br>(-0.13 to 0.04) | -0.03<br>(-0.16 to 0.02) | -0.04<br>(-0.16 to 0.02) |  | 0.00<br>(-0.13 to 0.08) | 0.02<br>(-0.14 to 0.18) |
| JHU (8M) | -0.01<br>(-0.10 to 0.03) | -0.03<br>(-0.12 to 0.01) | -0.04<br>(-0.12 to 0.02) | 0.00<br>(-0.08 to 0.13) |  | 0.02<br>(-0.08 to 0.17) |
| SR (8M) | -0.04<br>(-0.18 to 0.04) | -0.06<br>(-0.20 to 0.02) | -0.06<br>(-0.20 to 0.02) | -0.02<br>(-0.18 to 0.14) | -0.02<br>(-0.17 to 0.08) |  |

| Improvement in $\log_{10}$ error, IUPM = 2 | U. Pitt. | UCSD | JHU | SR | JHU (8M) | SR (8M) |
| --- | --- | --- | --- | --- | --- | --- |
| U. Pitt. |  | -0.01<br>(-0.03 to 0.01) | -0.01<br>(-0.04 to 0.04) | 0.01<br>(-0.03 to 0.07) | 0.02<br>(-0.02 to 0.07) | 0.03<br>(-0.02 to 0.13) |
| UCSD | 0.01<br>(-0.01 to 0.03) |  | -0.01<br>(-0.03 to 0.05) | 0.02<br>(-0.02 to 0.08) | 0.03<br>(-0.02 to 0.08) | 0.04<br>(-0.02 to 0.13) |
| JHU | 0.01<br>(-0.04 to 0.04) | 0.01<br>(-0.05 to 0.03) |  | 0.02<br>(-0.04 to 0.08) | 0.03<br>(-0.03 to 0.08) | 0.04<br>(-0.03 to 0.14) |
| SR | -0.01<br>(-0.07 to 0.03) | -0.02<br>(-0.08 to 0.02) | -0.02<br>(-0.08 to 0.04) |  | 0.01<br>(-0.06 to 0.08) | 0.02<br>(-0.04 to 0.11) |
| JHU (8M) | -0.02<br>(-0.07 to 0.02) | -0.03<br>(-0.08 to 0.02) | -0.03<br>(-0.08 to 0.03) | -0.01<br>(-0.08 to 0.06) |  | 0.01<br>(-0.06 to 0.11) |
| SR (8M) | -0.03<br>(-0.13 to 0.02) | -0.04<br>(-0.13 to 0.02) | -0.04<br>(-0.14 to 0.03) | -0.02<br>(-0.11 to 0.04) | -0.01<br>(-0.11 to 0.06) |  |

**S12 Table.** able Pairwise comparisons of assay accuracy for lower IUPMs, treating the JHU assay as gold standard (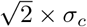 applied to other assays). Method and format match S10 Table.

| Improvement in $\log_{10}$ error, IUPM = 0.1 | U. Pitt. | UCSD | JHU | SR | JHU (8M) | SR (8M) |
| --- | --- | --- | --- | --- | --- | --- |
| U. Pitt. | | -0.36<br>(-0.53 to -0.20) | -0.37<br>(-0.73 to -0.01) | $+\infty$<br>(-0.30 to $+\infty$) | $+\infty$<br>(-0.55 to $+\infty$) | $+\infty$<br>(-0.11 to $+\infty$) |
| UCSD | 0.36<br>(0.20 to 0.53) | | -0.02<br>(-0.23 to 0.28) | $+\infty$<br>(0.06 to $+\infty$) | $+\infty$<br>(-0.11 to $+\infty$) | $+\infty$<br>(0.23 to $+\infty$) |
| JHU | 0.37<br>(0.01 to 0.73) | 0.02<br>(-0.28 to 0.23) | | $+\infty$<br>(0.03 to $+\infty$) | 0.60<br>(0.00 to $+\infty$) | $+\infty$<br>(0.18 to $+\infty$) |
| SR | $-\infty$<br>($-\infty$ to 0.30) | $-\infty$<br>($-\infty$ to -0.06) | $-\infty$<br>($-\infty$ to -0.03) | | 0.00<br>($-\infty$ to $+\infty$) | 0.00<br>(0.00 to $+\infty$) |
| JHU (8M) | $-\infty$<br>($-\infty$ to 0.55) | $-\infty$<br>($-\infty$ to 0.11) | -0.60<br>($-\infty$ to 0.00) | 0.00<br>($-\infty$ to $+\infty$) | | 0.00<br>(-0.08 to $+\infty$) |
| SR (8M) | $-\infty$<br>($-\infty$ to 0.11) | $-\infty$<br>($-\infty$ to -0.23) | $-\infty$<br>($-\infty$ to -0.18) | 0.00<br>($-\infty$ to 0.00) | 0.00<br>($-\infty$ to 0.08) | |

| Improvement in $\log_{10}$ error, IUPM = 0.2 | U. Pitt. | UCSD | JHU | SR | JHU (8M) | SR (8M) |
| --- | --- | --- | --- | --- | --- | --- |
| U. Pitt. | | -0.04<br>(-0.10 to 0.02) | -0.09<br>(-0.29 to 0.05) | 0.22<br>(-0.06 to $+\infty$) | -0.01<br>(-0.23 to $+\infty$) | 0.85<br>(-0.02 to $+\infty$) |
| UCSD | 0.04<br>(-0.02 to 0.10) | | -0.05<br>(-0.23 to 0.09) | 0.26<br>(-0.02 to $+\infty$) | 0.03<br>(-0.16 to $+\infty$) | 0.88<br>(0.02 to $+\infty$) |
| JHU | 0.09<br>(-0.05 to 0.29) | 0.05<br>(-0.09 to 0.23) | | 0.33<br>(0.01 to $+\infty$) | 0.11<br>(-0.02 to $+\infty$) | 1.03<br>(0.05 to $+\infty$) |
| SR | -0.22<br>($-\infty$ to 0.06) | -0.26<br>($-\infty$ to 0.02) | -0.33<br>($-\infty$ to -0.01) | | -0.19<br>($-\infty$ to $+\infty$) | 0.20<br>(-0.07 to $+\infty$) |
| JHU (8M) | 0.01<br>($-\infty$ to 0.23) | -0.03<br>($-\infty$ to 0.16) | -0.11<br>($-\infty$ to 0.02) | 0.19<br>($-\infty$ to $+\infty$) | | 0.55<br>(-0.11 to $+\infty$) |
| SR (8M) | -0.85<br>($-\infty$ to 0.02) | -0.88<br>($-\infty$ to -0.02) | -1.03<br>($-\infty$ to -0.05) | -0.20<br>($-\infty$ to 0.07) | -0.55<br>($-\infty$ to 0.11) | |

**S13 Table.** Table Pairwise comparisons of assay accuracy for higher IUPMs, treating the JHU assay as gold standard (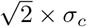 applied to other assays). Method and format match S10 Table.

| Improvement in $\log_{10}$ error, IUPM = 0.5 | U. Pitt. | UCSD | JHU | SR | JHU (8M) | SR (8M) |
| --- | --- | --- | --- | --- | --- | --- |
| U. Pitt. | | -0.04<br>(-0.10 to 0.02) | -0.13<br>(-0.29 to -0.04) | 0.02<br>(-0.08 to 0.41) | -0.06<br>(-0.23 to 0.08) | 0.06<br>(-0.09 to $+\infty$) |
| UCSD | 0.04<br>(-0.02 to 0.10) | | -0.08<br>(-0.24 to 0.02) | 0.07<br>(-0.03 to 0.42) | -0.02<br>(-0.20 to 0.14) | 0.10<br>(-0.03 to $+\infty$) |
| JHU | 0.13<br>(0.04 to 0.29) | 0.08<br>(-0.02 to 0.24) | | 0.15<br>(0.03 to 0.61) | 0.06<br>(-0.02 to 0.19) | 0.19<br>(0.02 to $+\infty$) |
| SR | -0.02<br>(-0.41 to 0.08) | -0.07<br>(-0.42 to 0.03) | -0.15<br>(-0.61 to -0.03) | | -0.09<br>(-0.54 to 0.08) | 0.05<br>(-0.15 to $+\infty$) |
| JHU (8M) | 0.06<br>(-0.08 to 0.23) | 0.02<br>(-0.14 to 0.20) | -0.06<br>(-0.19 to 0.02) | 0.09<br>(-0.08 to 0.54) | | 0.12<br>(-0.09 to $+\infty$) |
| SR (8M) | -0.06<br>($-\infty$ to 0.09) | -0.10<br>($-\infty$ to 0.03) | -0.19<br>($-\infty$ to -0.02) | -0.05<br>($-\infty$ to 0.15) | -0.12<br>($-\infty$ to 0.09) | |

| Improvement in $\log_{10}$ error, IUPM = 1 | U. Pitt. | UCSD | JHU | SR | JHU (8M) | SR (8M) |
| --- | --- | --- | --- | --- | --- | --- |
| U. Pitt. |  | -0.02<br>(-0.05 to 0.01) | -0.11<br>(-0.27 to -0.02) | 0.02<br>(-0.05 to 0.13) | -0.06<br>(-0.23 to 0.05) | 0.04<br>(-0.05 to 0.22) |
| UCSD | 0.02<br>(-0.01 to 0.05) |  | -0.09<br>(-0.26 to 0.00) | 0.03<br>(-0.03 to 0.14) | -0.05<br>(-0.22 to 0.07) | 0.05<br>(-0.03 to 0.24) |
| JHU | 0.11<br>(0.02 to 0.27) | 0.09<br>(0.00 to 0.26) |  | 0.13<br>(0.03 to 0.32) | 0.05<br>(-0.01 to 0.12) | 0.15<br>(0.03 to 0.41) |
| SR | -0.02<br>(-0.13 to 0.05) | -0.03<br>(-0.14 to 0.03) | -0.13<br>(-0.32 to -0.03) |  | -0.08<br>(-0.28 to 0.04) | 0.02<br>(-0.12 to 0.20) |
| JHU (8M) | 0.06<br>(-0.05 to 0.23) | 0.05<br>(-0.07 to 0.22) | -0.05<br>(-0.12 to 0.01) | 0.08<br>(-0.04 to 0.28) |  | 0.10<br>(-0.03 to 0.36) |
| SR (8M) | -0.04<br>(-0.22 to 0.05) | -0.05<br>(-0.24 to 0.03) | -0.15<br>(-0.41 to -0.03) | -0.02<br>(-0.20 to 0.12) | -0.10<br>(-0.36 to 0.03) |  |

| Improvement in $\log_{10}$ error, IUPM = 2 | U. Pitt. | UCSD | JHU | SR | JHU (8M) | SR (8M) |
| --- | --- | --- | --- | --- | --- | --- |
| U. Pitt. |  | -0.01<br>(-0.03 to 0.02) | -0.10<br>(-0.26 to -0.01) | 0.01<br>(-0.03 to 0.08) | -0.06<br>(-0.23 to 0.03) | 0.03<br>(-0.02 to 0.12) |
| UCSD | 0.01<br>(-0.02 to 0.03) |  | -0.09<br>(-0.26 to -0.01) | 0.02<br>(-0.03 to 0.09) | -0.05<br>(-0.22 to 0.04) | 0.03<br>(-0.02 to 0.12) |
| JHU | 0.10<br>(0.01 to 0.26) | 0.09<br>(0.01 to 0.26) |  | 0.12<br>(0.01 to 0.30) | 0.04<br>(-0.01 to 0.09) | 0.13<br>(0.03 to 0.32) |
| SR | -0.01<br>(-0.08 to 0.03) | -0.02<br>(-0.09 to 0.03) | -0.12<br>(-0.30 to -0.01) |  | -0.07<br>(-0.26 to 0.03) | 0.01<br>(-0.05 to 0.10) |
| JHU (8M) | 0.06<br>(-0.03 to 0.23) | 0.05<br>(-0.04 to 0.22) | -0.04<br>(-0.09 to 0.01) | 0.07<br>(-0.03 to 0.26) |  | 0.09<br>(-0.02 to 0.28) |
| SR (8M) | -0.03<br>(-0.12 to 0.02) | -0.03<br>(-0.12 to 0.02) | -0.13<br>(-0.32 to -0.03) | -0.01<br>(-0.10 to 0.05) | -0.09<br>(-0.28 to 0.02) |  |

**S14 Table.** Table Simulated results of latency reduction trials, maximum likelihood estimation to compare pre- and post-treatment data.

|  | JHU protocol |  |  |  |
| --- | --- | --- | --- | --- |
| | Weaker therapy (3x),<br>$N = 12$ per arm | | Stronger therapy (10x),<br>$N = 6$, no control arm | |
|  | Unbatched | Batched | Unbatched | Batched |
| Median estimated fold-reduction | 3.13 | 3.06 | 10.67 | 9.91 |
| Median bias | +4% | +2% | +7% | −1% |
| Median $\log_{10}$ absolute error | 0.10 | 0.08 | 0.23 | 0.21 |
| Accuracy improvement from batching (% reduction in abs. error) | 20% |  | 9% |  |
| Coverage of 95% CI | 88.4% | 87.2% | 91.0% | 90.4% |
| Power ($p < 0.05$) | 90.5% | 98.6% | 77.5% | 79.2% |
| Power improvement from batching (percentage point increase) | 8% |  | 2% |  |

|  | UCSD protocol |  |  |  |
| --- | --- | --- | --- | --- |
| | Weaker therapy (3x),<br>$N = 12$ per arm | | Stronger therapy (10x),<br>$N = 6$, no control arm | |
|  | Unbatched | Batched | Unbatched | Batched |
| Median estimated fold-reduction | 3.12 | 3.03 | 10.52 | 9.71 |
| Median bias | +4% | +1% | +5% | −3% |
| Median $\log_{10}$ absolute error | 0.11 | 0.08 | 0.21 | 0.18 |
| Accuracy improvement from batching (% reduction in abs. error) | 27% |  | 14% |  |
| Coverage of 95% CI | 88.4% | 87.3% | 90.7% | 89.3% |
| Power ($p < 0.05$) | 90.5% | 99.4% | 83.2% | 90.9% |
| Power improvement from batching (percentage point increase) | 9% |  | 8% |  |

**S1 Fig.**
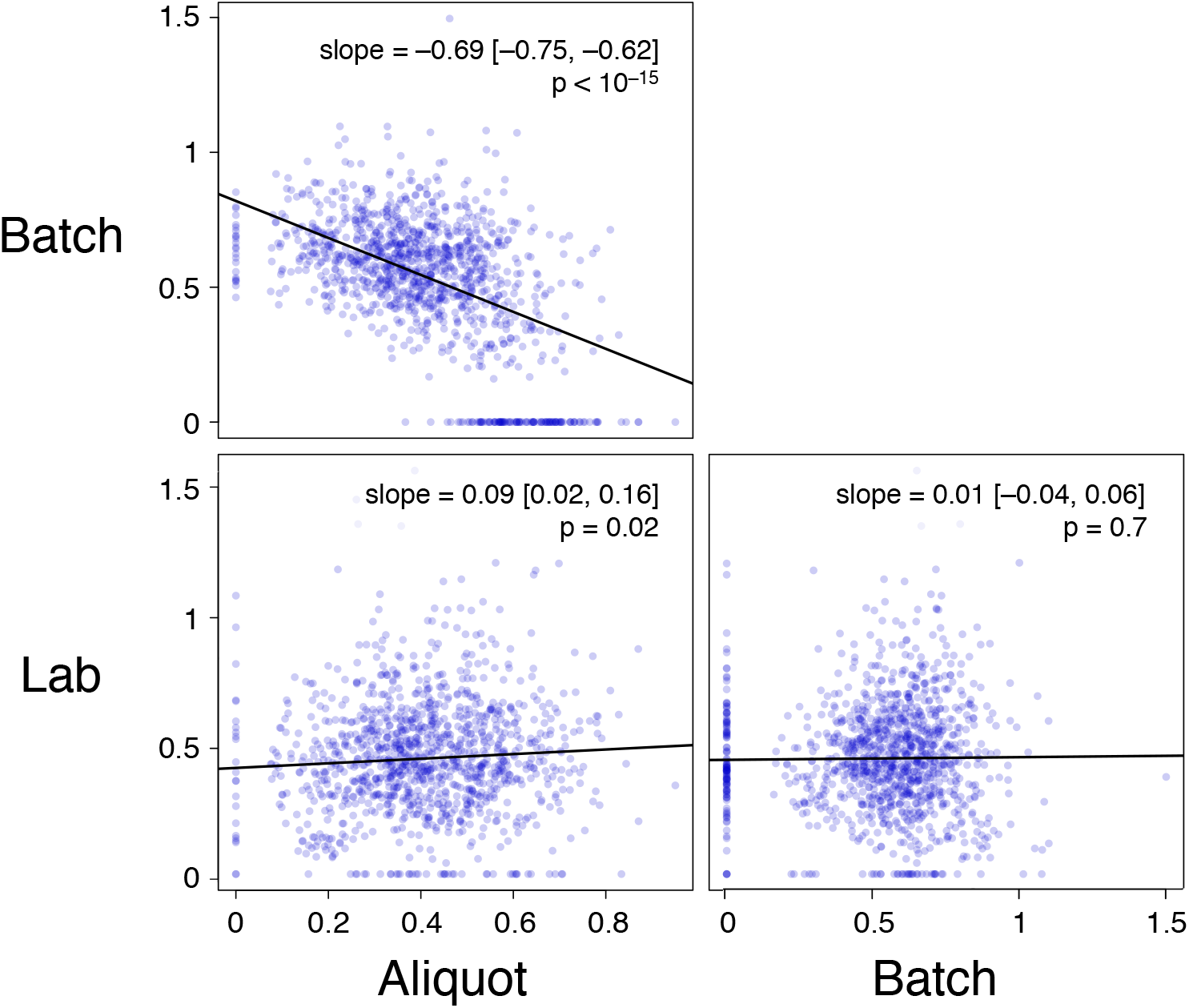
Scatterplot of estimates of excess variation at the aliquot, batch, and lab levels, expressed as the natural log of fold-change (parameters *σ_a_, σ_b_, σ_c_*). Panels show robust linear regression and Wald test p-values (null model = zero slope) of 1000 samples from the ensemble posterior.

**S2 Fig.**
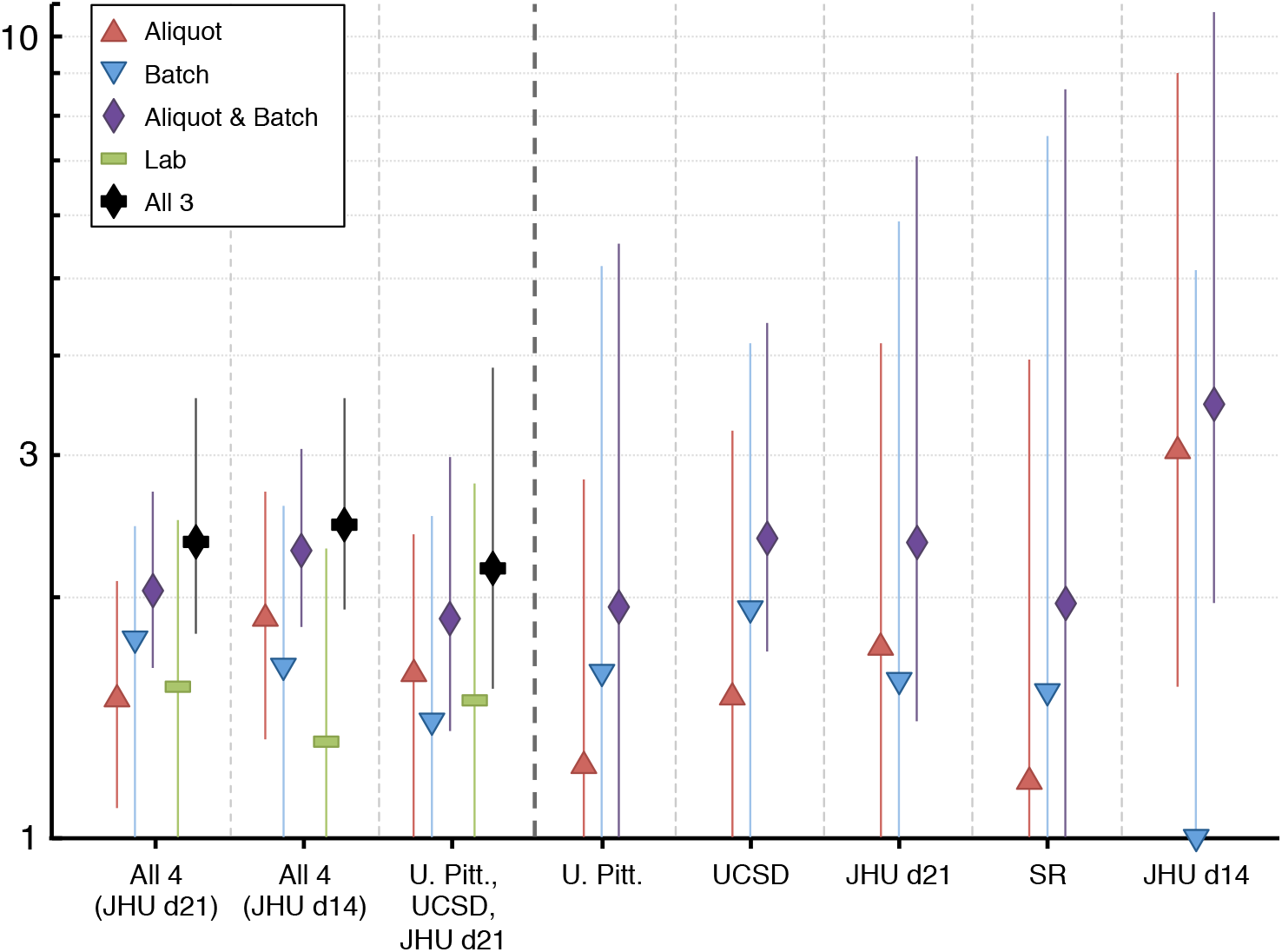
Sensitivity analysis of of aliquot-level, batch-level, and lab-level variation.

**S3 Fig.**
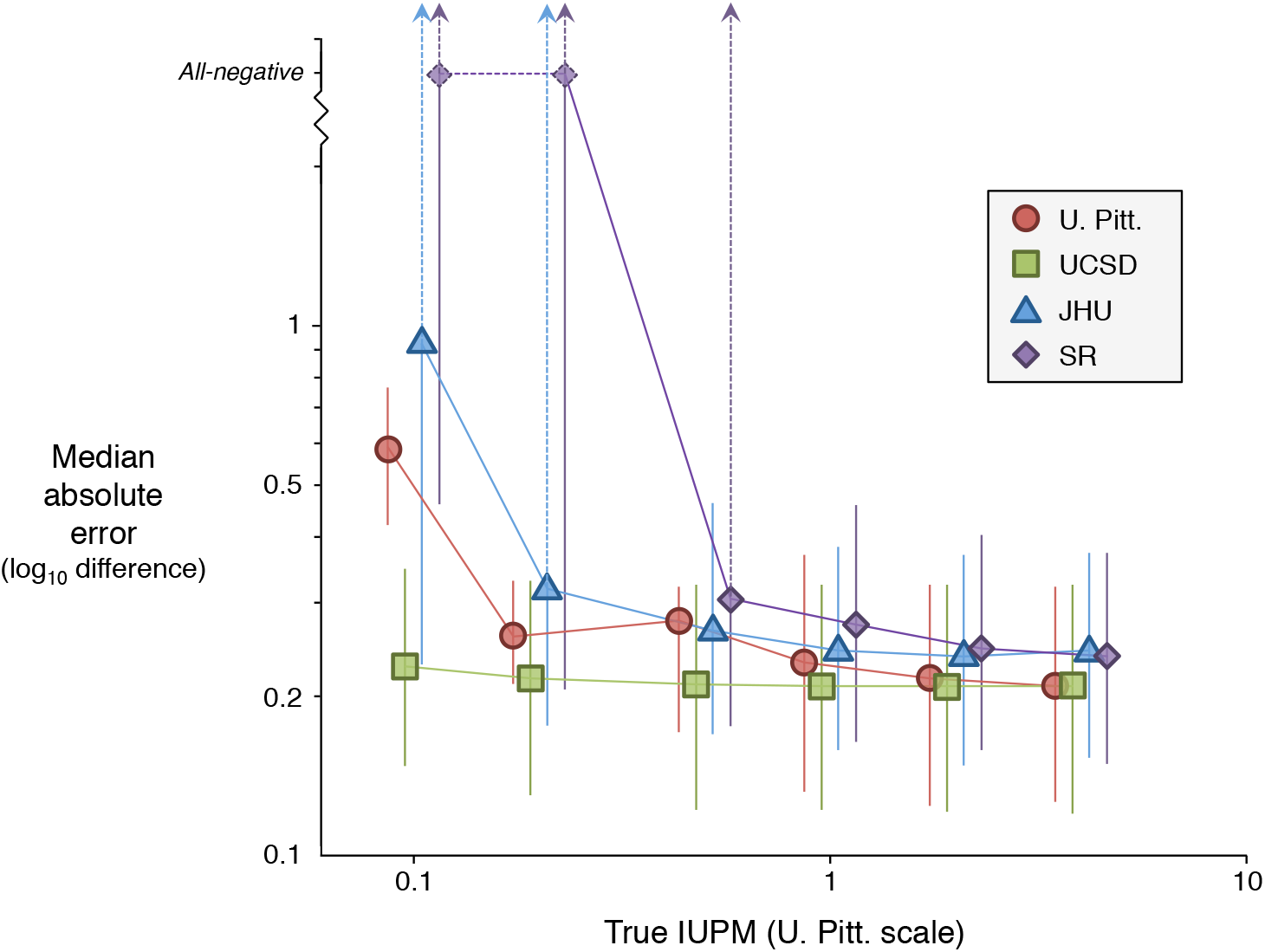
Accuracy of assays used in the experimental study, with JHU and SR assays modified to use the same number of input cells as U. Pitt. and UCSD. Each assay is measured against a consensus standard, appropriately scaled by *β_l_* for that assay. “All-negative” on the *y*-axis represents infinite error on the log scale, which occurs when the maximum likelihood estimate of IUPM is zero.

### S1 Appendix. Stan model code

Complete model specifications are provided in the Stan programming language (version 2.12) as supplementary text files. Each filename has a binary code, indicating presence/absence of aliquot and batch effects (for single-lab models) or presence/absence of aliquot, batch, and lab effects (for multi-lab models). In the code, `SourcelD` refers to participant index *i*, `UniqAliquot` refers to concatenated participant-aliquot index *ij*, `UniqBatch` refers to concatenated batch-lab index *kl*, `AssaySourcelD` refers to concatenated participant-lab index *il*, `Assay` refers to lab index *l*, and `Stored` refers to cryopreservation indicator *S_ij_*. Note that all effects are represented as logit values, ensuring that all frequencies remain between 0 and 1 as MCMC explores the parameter space.

## Acknowledgments

A Collaboration for AIDS Vaccine Discovery (CAVD) grant from the Bill & Melinda Gates Foundation (Grant ID: 38619) supports the RAVEN Study Group. DISR was supported by amfAR Mathilde Krim Fellowship # 109511-61-RKRL. Work performed at SR was supported in whole by federal funds from the NIH/NIAID/DAIDS under contract HHSN272201500017C entitled “Quantitative Viral Outgrowth Assay (QVOA) Service Resource.” Members of the RAVEN Study Group: *Steering Committee*: Michael P. Busch, Silvija Staprans, John Mellors, Doug Richman, Steve Deeks, Peter Bacchetti, Joe Fitzgibbon. *Central Lab Staff*: Mars Stone, Sheila Keating, Sonia Bakkour, Xutao Deng. *Key Collaborators*: Ron Bosch, Rebecca Ho, Robert Siliciano, Janet Siliciano, Nicolas Chomont, Daniel Rosenbloom, Roger Ptak, Sulggi Lee.

